# Artificial intelligence networks combining histopathology and machine learning can extract axon pathology in autism spectrum disorder

**DOI:** 10.1101/2024.10.25.620308

**Authors:** Arash Yazdanbakhsh, Kim Dang, Kelvin Kuang, Tingru Lian, Xuefeng Liu, Songlin Xie, Basilis Zikopoulos

**Affiliations:** Computational Neuroscience and Vision Laboratory, Department of Psychological and Brain Sciences, Boston University, Boston, MA, 02215 United States; Graduate Program for Neuroscience, Boston University, Boston, MA, United States; Center for Systems Neuroscience, Boston University, Boston, MA, United States; Human Systems Neuroscience Laboratory, Department of Health Sciences, Program in Human Physiology, Boston University, Boston, MA, 02215 United States; Department of Anatomy & Neurobiology, Boston University School of Medicine, Boston, MA, United States

**Keywords:** white matter, anterior cingulate cortex, short-range pathways, long-range pathways, deep neural network, convolutional neural network

## Abstract

Axon features that underlie the structural and functional organization of cortical pathways have distinct patterns in the brains of neurotypical controls (CTR) compared to individuals with Autism Spectrum Disorder (ASD). However, detailed axon study demands labor-intensive surveys and time-consuming analysis of microscopic sections from post-mortem human brain tissue, making it challenging to systematically examine large regions of the brain. To address these challenges, we developed an approach that uses machine learning to automatically classify microscopic sections from ASD and CTR brains, while also considering different white matter regions: superficial white matter (SWM), which contains a majority of axons that connect nearby cortical areas, and deep white matter (DWM), which is comprised exclusively by axons that participate in long-range pathways. The result was a deep neural network that can successfully classify the white matter below the anterior cingulate cortex (ACC) of ASD and CTR groups with 98% accuracy, while also distinguishing between DWM and SWM pathway composition with high average accuracy, up to 80%. Multidimensional scaling analysis and sensitivity maps further underscored the reliability of ASD vs CTR classification, based on the consistency of axon pathology, while highlighting the important role of white matter location that constrains pathway dysfunction, based on several shared anatomical markers. Large datasets that can be used to expand training, validation, and testing of this network have the potential to automate high-resolution microscopic analysis of post-mortem brain tissue, so that it can be used to systematically study white matter across brain regions in health and disease.

**One Sentence Statement:** Histopathology-trained AI can identify ASD network disruptions and guide development of diagnostics and targeted therapeutics.

## Introduction

Neural communications and connections are altered in Autism Spectrum Disorder (ASD) and several studies have pinpointed changes in axons in few frontal, temporal, and callosal regions, as a core underlying pathology (*1–8*). In these studies, the most observed axon pathology involves an increase in the relative density of thin axons accompanied by a parallel decrease in the density of thick axons, however, excessive branching due to expression of growth axon proteins, thinning of the myelin, changes in the inner/outer diameter ratio (g-ratio), and increased variability of the trajectory of axons have also been reported.

These structural and molecular changes of individual axons underlie conduction speed and pathway strength, and ultimately determine the physiology and fidelity of signal transmission, and the integrity of neural communications (*9–12*). In addition, the relative position and size of axons in the white matter can be used as an indicator of their termination in nearby or distant brain areas. The superficial white matter (SWM) mainly includes short-range connections whereas, the deep white matter (DWM) includes long range excitatory pathways, with axons that are significantly thicker than axons found in the SWM, (*1, 13, 14*). Therefore, systematic study of the white matter can reveal key structural and functional principles of the organization of brain pathways and mechanisms of disruption in ASD. However, high-resolution quantitative examination of features of individual axons requires extensive study by experienced anatomists. Such analyses are labor-intensive and time-consuming, rendering this approach not optimal for large-scale studies that can help identify core ASD network status and likely mechanisms of disruption in communication.

Machine learning may offer an alternative approach for the classification of high-resolution microscopic images of myelinated axons in white matter pathways that link nearby and distant cortical and subcortical areas in neurotypical controls and individuals with ASD. Deep neural networks (DNN) are tools employed in machine learning that have performed incredibly well with image classification (*15*). Moreover, DNN and transfer learning are increasingly used to classify medical images, showing the strengths of such an approach for spatial feature detection [(*16–19*), reviewed in (*20, 21*)]. Among the various deep neural networks, GoogLeNet is a convolutional neural network that is established as a good prototype with sufficient power for small-scale datasets (*15*). Therefore, in this study we used GoogLeNet to develop a template for customization of a deep neural network through transfer learning that would be capable of distinguishing high-resolution microscopic images of myelinated axons in white matter pathways that link nearby and distant brain areas in neurotypical controls and individuals with ASD. We focused on the white matter below the anterior cingulate cortex (ACC), that has a key role in attention, social interactions, emotions, and executive control, processes that are affected in autism (*22–25*). The ACC is one of the brain regions most consistently and severely affected in ASD, exhibiting hyperactivity during response monitoring and social target detection (*26, 27*) and desynchronized activity during working memory tasks (*28*), likely due to local over-connectivity and long distance disconnection of ACC pathways (*29–33*). To develop and optimize initial machine learning algorithms we used a large dataset of light and electron microscopy (EM) images of white matter axons from short- and long-range ACC cortical pathways in control and ASD groups of adults from studies we published since 2010 that have shown excessive branching and thinning of axons linking ACC with nearby areas in the SWM and fewer thick axons traveling long distances in the DWM in ASD (*1–3, 34*).

Classification of pathology and white matter depth by our trained network reached high levels of reliability. Quantification and visualization of regions highlighted as crucial for correct and incorrect network classification reaffirmed previous findings and provided new insights, revealing novel features and cues that were not apparent in histopathological analysis and underlie ASD heterogeneity. Our proposed method can be generalized and applied to detect axonal and pathway feature differences across brain areas in the neurotypical brain and in psychiatric disorders with underlying pathology in brain network connectivity.

## Materials and Methods

### Tissue Processing and Dataset Selection

We used two datasets, consisting of thousands of digital photomicrographs from post-mortem human brain sections of the white matter below ACC that were processed and imaged in the Human Systems Neuroscience Laboratory at Boston University (B. Zikopoulos, PI), as described in previous studies (*1–3, 34*). The first dataset consisted of nanometer-scale electron microscopy (EM) images, and the second dataset included micrometer-scale brightfield optical microscopy images stained with the Nissl stain toluidine blue (Tol-Blue).

The brief description below summarizes previously reported relevant protocols for tissue selection, processing, and imaging (*1–3, 34*). We obtained age- and gender-matched post-mortem brain tissue that was immersion fixed in 10% formalin, from 12 individuals, 7 neurotypical controls (CTR) and 5 with ASD (Table 1) from the Autism Tissue Program, the Harvard Brain Tissue Resource Center, the Institute for Basic Research in Developmental Disabilities, the University of Maryland Brain and Tissue Bank, the National Disease Research Interchange (NDRI), Anatomy Gifts Registry, and Autism BrainNet. The diagnosis of autism was based on the Autism Diagnostic Interview-Revised (ADI-R). Clinical characteristics, including autism diagnostic interview scores, and other data are summarized in Table 1. Cases were matched for the most part, based on tissue availability, however a larger sample of control cases with a wider age-range that included older individuals was used for initial training purposes. The study was approved by the Institutional Review Board of Boston University.

**Table 1.** Demographics, indices, and scores for each individual case. PMI: post-mortem interval. Generally, shorter PMI correlates with better tissue quality for histopathological studies however, we obtained well-preserved and appropriately stored tissue with a short PMI (on average 31.6 h or 25.5 h, when excluding one case with PMI of 99 h), despite limited tissue availability. Our sample included one ASD case with a PMI of 99 h; however, the mean PMI overall was relatively low and comparable to the mean PMI of cases used in similar studies. Signs of post-mortem autolysis were minimal in myelinated axons and the white matter and were restricted to minor dissociation of the myelin sheath or small inclusions in the axolemma in some thick axons, and the typical ‘fried egg’ appearance of oligodendrocytes in *post-mortem* tissue, with a seemingly intact nucleus positioned in a bloated, but empty cytoplasm. ASD_x_: Cases from ASD adults, CTR_x_: Cases from the control group, *: exclusive to EM images, not in training for Tol-Blue images; ** Asphyxiation or respiratory problems may result in suboptimal tissue quality; *** The number next to each criterion of the ADI-R is the cut-off score of that criterion. An autism diagnosis is indicated when scores in all three behavioral areas meet or exceed the specified minimum cutoff scores. The three criteria are: social interaction (S), communication and language (C): verbal (V) and nonverbal (NV), restrictive and repetitive behaviors (R).

| Case Number | Age (y) | Sex | PMI (h) | Cause of Death | ADI-R*** |  |  |
| --- | --- | --- | --- | --- | --- | --- | --- |
|  |  |  |  |  | S(10) | C(8V, 7NV) | R(3) |
| ASD<sub>1</sub> | 30 | M | 16 | Congestive heart failure | 26 | 22(V) | 12 |
| ASD<sub>2</sub> | 30 | M | 20 | Myocardial infarction | 22 | 12(NV) | 12 |
| ASD<sub>3</sub> | 44 | M | 31 | Acute myocardial infarction | 26 | 18(V), 13(NV) | 6 |
| ASD<sub>4</sub> | 40 | F | 33 | Respiratory arrest** | 12 | 14(V) | 8 |
| ASD<sub>5</sub> | 31 | M | 99 | Shooting | 18 | 14(V) | 6 |
| CTR<sub>1</sub> | 42 | M | 18 | Myocardial infarction | NA | NA | NA |
| CTR<sub>2</sub> | 36 | F | 18 | Unknown | NA | NA | NA |
| CTR<sub>3</sub> | 36 | M | 20 | Myocardial infarction | NA | NA | NA |
| CTR<sub>4</sub> | 67 | M | 30 | Pancreatic cancer | NA | NA | NA |
| CTR<sub>5</sub> | 58 | F | 30 | Pancreatic cancer | NA | NA | NA |
| CTR*<sub>6</sub> | 83 | M | 24 | Unknown | NA | NA | NA |
| CTR<sub>7</sub> | 30 | M | 41 | Unknown | NA | NA | NA |

We used coronal ACC tissue blocks, matched based on the human brain atlas (*35*) and (*36*), and additional cytoarchitectonic studies of human prefrontal and cingulate cortex (*37–40*). We postfixed tissue slabs in 2% paraformaldehyde and 2.5% glutaraldehyde, in 0.1 M phosphate buffer (PB, pH: 7.4) for 2–4 days at 4 °C. To preserve the ultrastructure until processing, we cryoprotected tissue blocks in gradually-increasing sucrose solutions (10-25%) and then immersed them in anti-freeze solution (30% ethylene glycol, 30% glycerol, 40% 0.05 M PB, pH: 7.4 with 0.05% azide) before storing at − 20 °C. We then rinsed tissue blocks in 0.1 M PB and cut them coronally at 50-μm-thick sections on a vibratome (Pelco, series 1000). Some blocks were frozen in − 70 °C isopentane and cut in a cryostat (CM 1500, Leica), or a sliding microtome (AO), equipped with a PhysiTemp freezing platform, in the coronal plane at 20–50 µm in 10 series of free-floating sections.

### Electron Microscopy Processing and Imaging

For electron microscopy (EM) processing, cutting, and staining, we used adjacent series of 50-μm-thick coronal sections and stained with a high-contrast method. We confirmed the presence of ACC cortical gray matter in adjacent sections that were stained with Nissl. We rinsed sections in 0.1 M PB and postfixed them in a variable wattage microwave oven (Biowave, Pelco) with 6% glutaraldehyde at 100–150 W. Under a dissecting microscope, we cut small regions of sections containing the superficial or deep parts of the white matter below ACC. White matter regions of interest were rinsed and incubated in the following series of solutions: 0.1 M cacodylate buffer, then postfixed with intermediate water rinses first in 0.1% tannic acid, followed by 1% osmium tetroxide with 1.5% potassium ferrocyanide, then TCH aqueous solution (0.1 g of thiocarbohydrazide), and finally 2% osmium tetroxide. Sections were washed in water, stained overnight with 1% uranyl acetate followed by lead aspartate and dehydrated in an ascending series of alcohols. We used propylene oxide to clear tissue sections before embedding in araldite or LX112 resin at 60 °C and then in Aclar film for storage. We prepared resin blocks and cut serial semithin (1 μm) or ultrathin (50-100 nm) coronal sections with a diamond knife (Diatome, Fort Washington, PA), using an ultramicrotome (Ultracut; Leica, Wein, Austria). EM processing rendered lipids in the myelin sheath and in membranes electron-dense and they appeared dark (*41*). We collected ultrathin sections on single slot pioloform-coated grids to view and acquire images with a scanning electron microscope (Zeiss Gemini 300 with STEM detector and Atlas 5 software modules), at high-resolution (10 nm/pixel), as described (*1–4, 6*).

### Light Microscopy Processing and Imaging

For the additional Nissl staining of semithin sections, we used Toluidine Blue, which stains all cells and the neuropil. We prepared 1% aqueous solution with toluidine blue powder (T3260, Sigma-Aldrich) in distilled water, filtered and mixed with sodium borate (1:1; S9640, Sigma-Aldrich), filtered again and diluted 1:1 with 70% ethanol. Semithin sections floating in water were mounted on gelatin-coated slides and placed on a heater plate until the water evaporated. Sections were then covered with the final toluidine blue solution for ≤1 min, rinsed with water, differentiated with 30-70% ethanol and rinsed with water, before coverslipping with Entellan. We photographed sections using an optical microscope (Olympus optical microscope, BX51) with a CCD camera (Olympus DP70) connected to a personal computer with a commercial imaging system (DP Controller). Photographs were taken with the 100× objective (Olympus UPlanFL N 100×/1.30 Oil Iris ∞/0.17/FN26.5) using oil immersion (Immersol^TM^, 518F, Zeiss, Oberkochen, Germany).

Images in each dataset (EM and Tol-Blue) were labeled based on the population type from which the sections were obtained: neurotypical control group (CTR) versus adults with ASD, and further divided according to the location of origin within the white matter: superficial white matter (SWM) versus deep white matter (DWM). The resulting four classes were labeled as “Control Deep White Matter” (CTR DWM), “Control Superficial White Matter” (CTR SWM), “Autism Spectrum Disorder Deep White Matter” (ASD DWM), and “Autism Spectrum Disorder Superficial White Matter” (ASD SWM). The EM dataset consisted of images at 2500 x 2500 pixels (10nm/ pixel), that covered and sampled an average white matter surface area of 0.3 mm^2^/case in each of the groups (ASD DWM, ASD SWM, CTR DWM, CTR SWM). The optical microscopy dataset consisted of images with an average dimension of 4080 x 3072 pixels (42.78 nm/pixel), that covered and sampled an average white matter surface area of 1.6 mm^2^/case in each of the groups (ASD DWM, ASD SWM, CTR DWM, CTR SWM).

### Image Preprocessing Methods

As network training typically benefits from a large dataset, we considered two image pre-processing methods that would increase the size of the dataset: the N x N window-tiling method, and the sliding window method. Both methods were applied to determine the optimal sub-image size for best network accuracy and implemented by nested “for” loops in MATLAB 2023a. The N x N window-tiling method involved cutting the original image into smaller sub-images by splitting each dimension by N, creating N^2^ images. For example, if N = 2, one original image would generate 4 sub-images. By considering different N values, each classification accuracy was evaluated and compared with others, and the best-performing N value with its corresponding sub-image size was determined. After implementation of the window-tiling method, for EM we had an average of 2,500 images (ASD) and 2,000 images (CTR), while there were 7,380 images per class for the Tol-Blue slices.

The second image pre-processing method we considered was the sliding window method. Similarly to the sub-imaging method, the larger image would be divided into smaller sub-images, but a sliding window was applied, resulting in overlaps within the sub-images (Figure 1). For the electron microscopy dataset, the sliding window moved 500 pixels (1/4ths the sub-image size) for each sub-image in both the x- and y-axes. For the optical microscopy dataset, the sliding window moved 89 pixels (1/5ths the sub-image size) for each sub-image in both the x- and y-axes.

**Figure 1:**
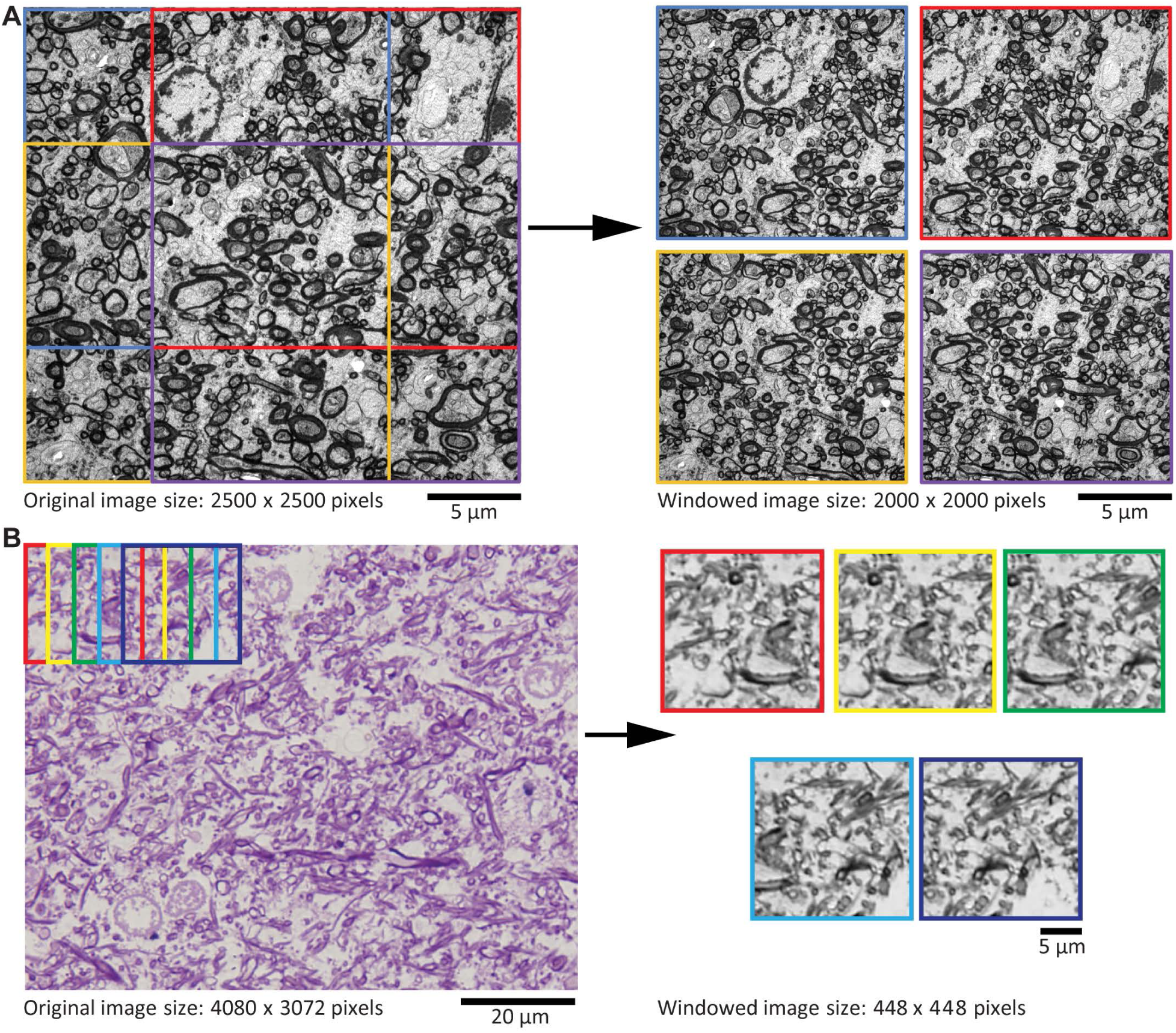
Specifications and preprocessing of microscopic section images. **A.** An example of a source EM section image, that can be digitally cropped to generate subsections to be used as DNN input; note the color correspondence of square outlines of cropped areas before and after the arrow to visualize subsections from the source section. **B.** An illustration of the sliding window method used for Tol-Blue images; note the stride of windows (sequence of colored square outlines) and the resultant sections to be input to DNN. Again, the square outline color correspondence before and after the arrow shows corresponding subsections.

In addition, for the Tol-Blue optical microscopy dataset, the images were first converted to grayscale to remove any background features that the color of the dye may have introduced. Due to the varying contrasts within the slices, contrast-limited adaptive histogram equalization was also applied to create a more uniform dataset by utilizing Image Processing and Computer Vision, and Statistics & Machine Learning toolboxes of MATLAB 2023a. The images were also normalized to the values between [0, 1] instead of [0, 255] for training.

To avoid overfitting in the trained network, we employed a data splitting method that would ensure a complete separation across cases between the training and testing datasets. The EM images and the optical microscopy images were separated differently due to their variances in case and image availability. Specifically, for the EM images we used 2 different methods of data distribution between training, validation, and testing datasets (Table 2). The first method separated the training/validation data from the testing dataset, with one case (∼20% of the dataset) going into the testing dataset, while the rest of the cases were split 7:3 to the training/validation set. The second method separated the validation/testing dataset from the training dataset, with one case being split 50:50 to the validation/testing dataset, while the rest of the cases went to the training set.

**Table 2:**
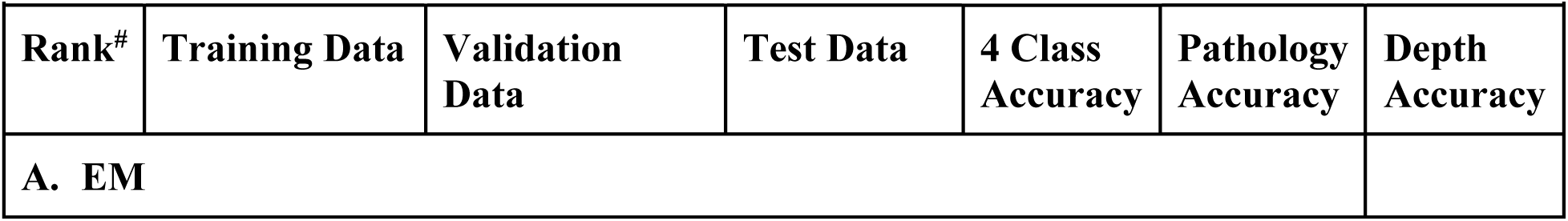
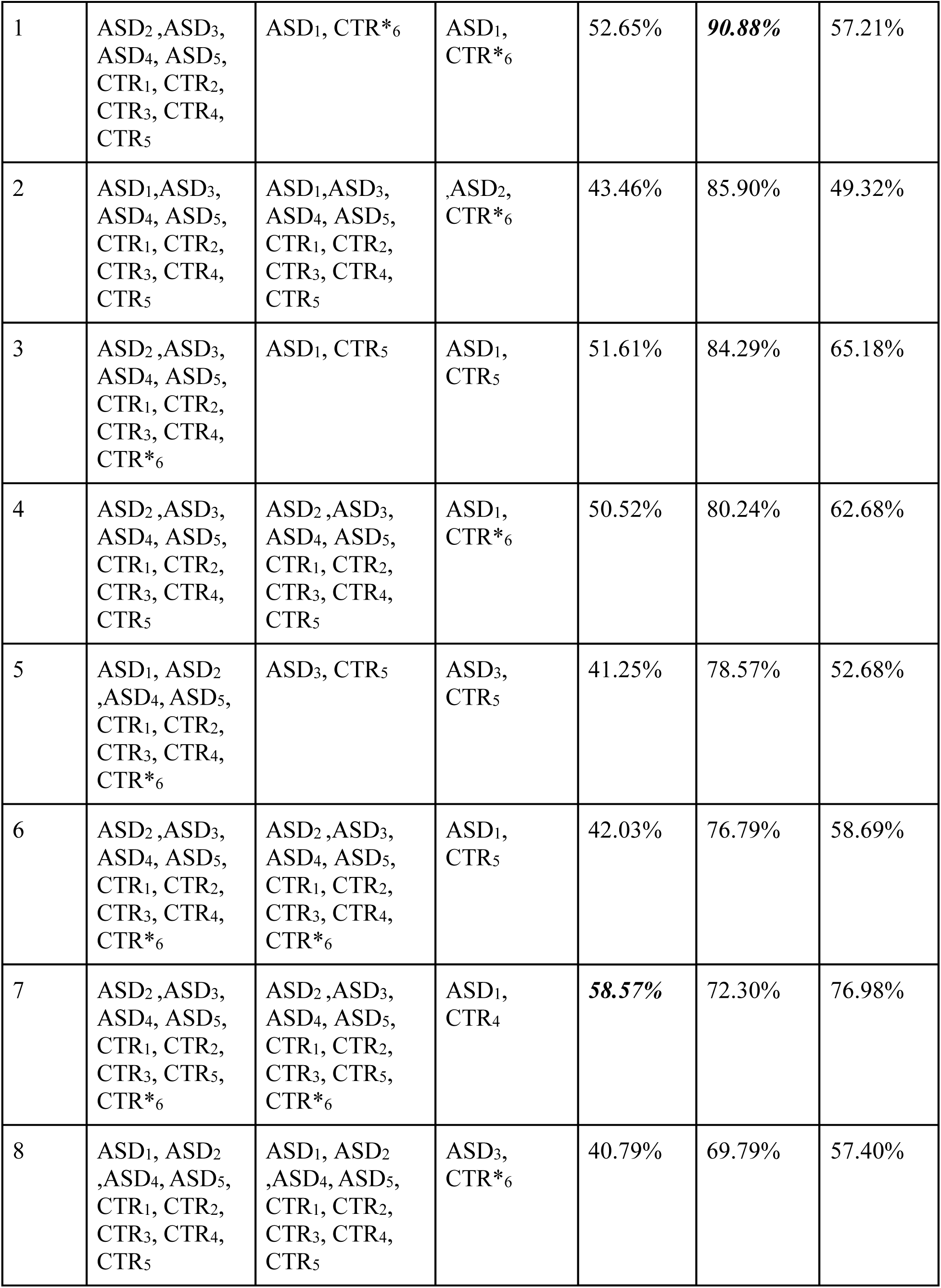
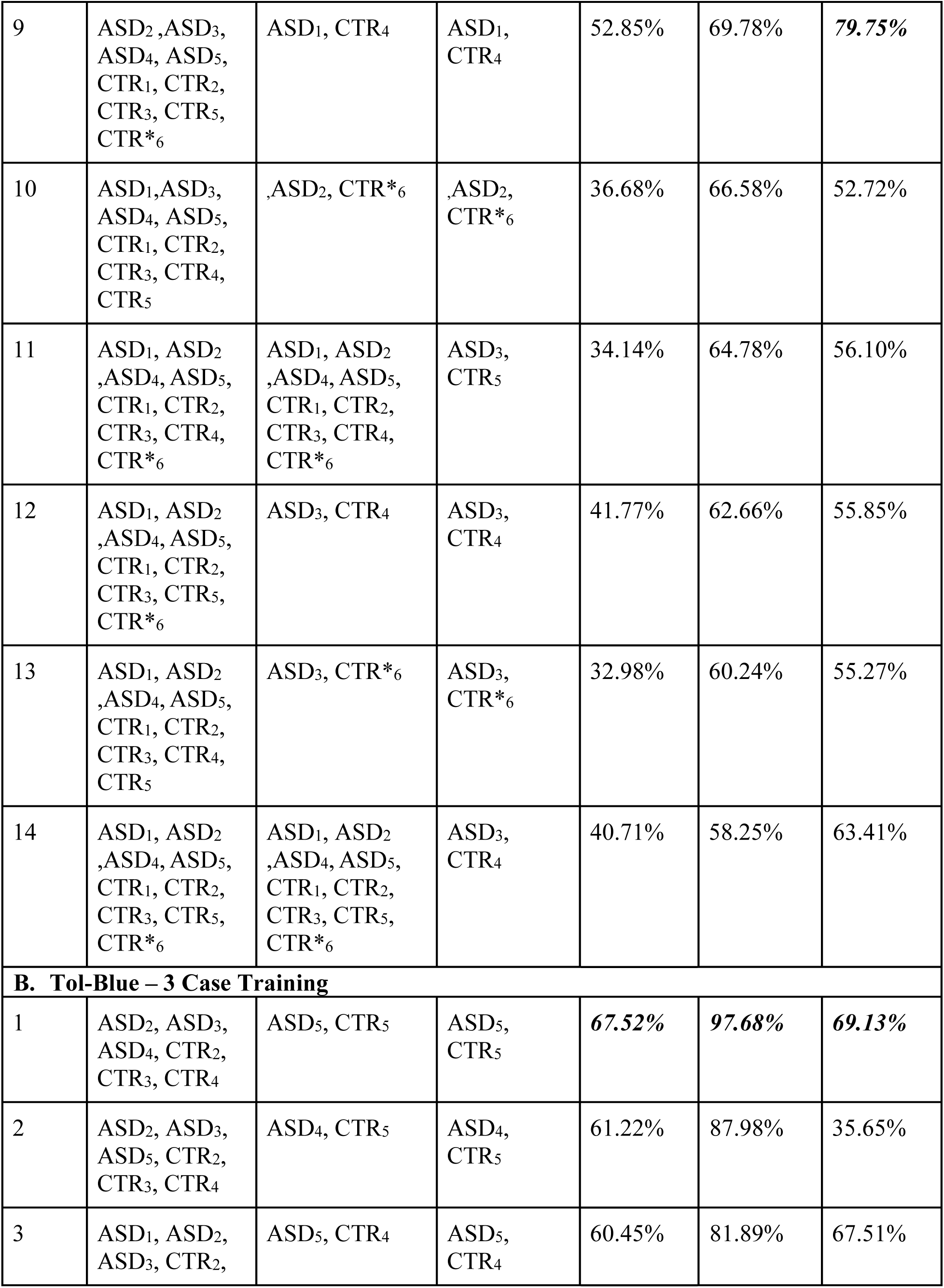
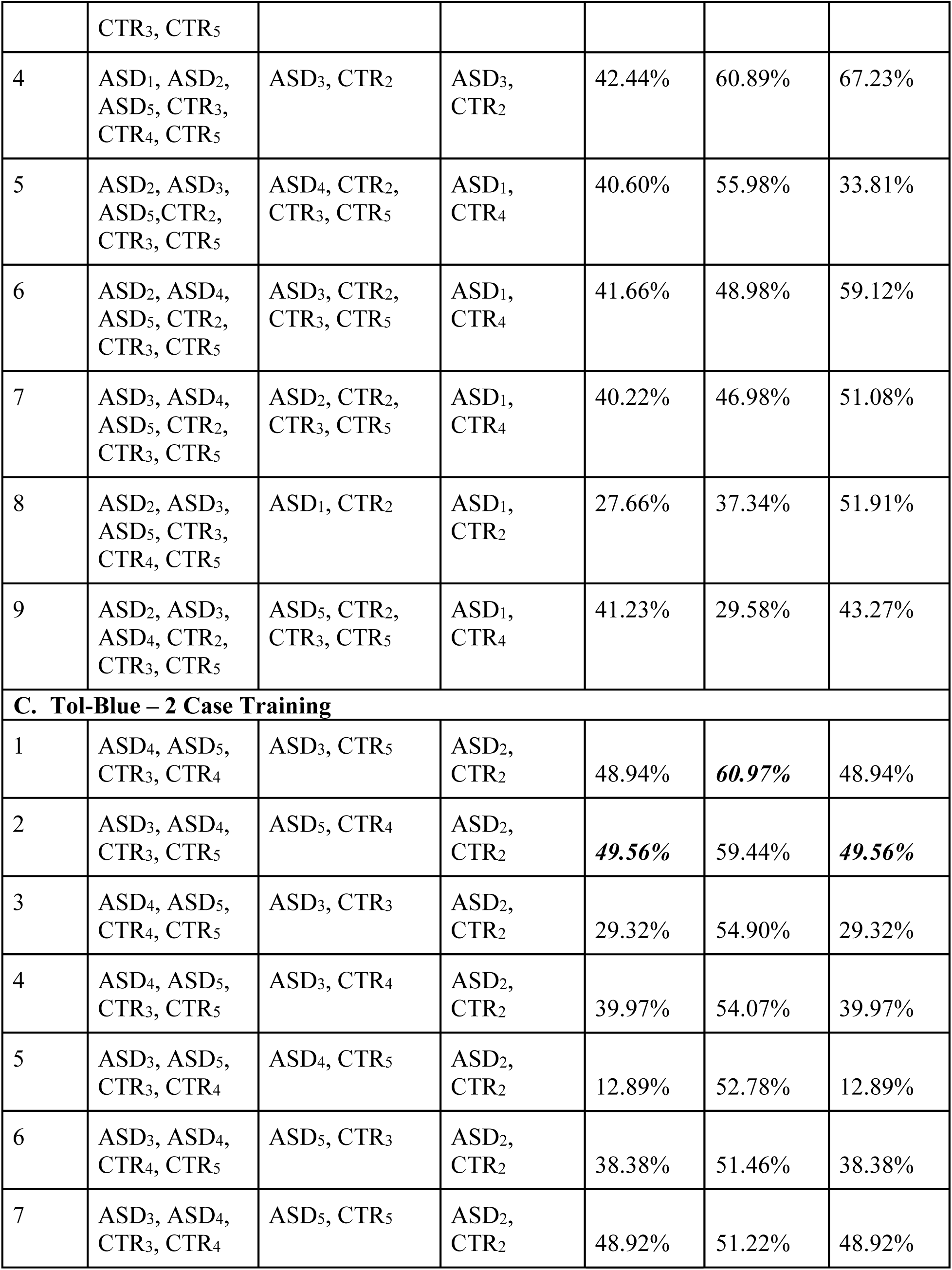
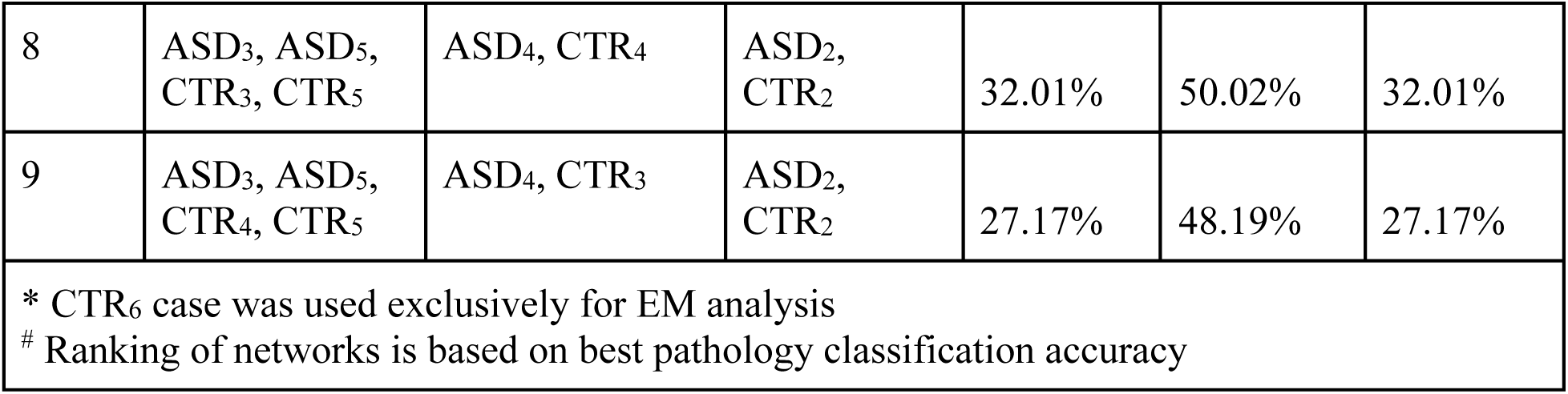
Rank-based training on different case combinations. Ranking is based on pathology classification accuracy (descending). **A.** Results for EM images. **B.** Results for 3 case training (3 CTR and 3 ASD) on Tol-Blue images. **C.** Results for 2 case training (2 CTR and 2 ASD) on Tol-Blue images. Our exhaustive training and testing with different combinations of non-overlapping cases, showed that with small datasets, in this case limited number of cases and images, the best approach was to train separate models for optimal classification of pathology only, or white matter depth only, or combinations of these (best classification accuracies in ***italicized bold*** font). Note that there was no overlap of cases and images between training and testing. In addition, there was no overlap of images between training and validation, even when training and validation sets came from the same cases (e.g., EM ranked-based training sessions 2, 4, 6, 7 etc.).

### Model Training, Validation, and Testing

To increase generalizability and reliability of the model we ensured that each training and testing set included images from different cases and there were no shared sub-images in either dataset. Therefore, we ensured that there was no overlap of cases and images between training and testing. In addition, there was no overlap of images between training and validation, even when training and validation sets came from the same cases. We addressed the variability in the number of images across cases by employing both undersampling and oversampling techniques to balance the training dataset for each class. We utilized undersampling by identifying the class with the fewest images and limiting the training set for each class to that minimum number through random selection. Conversely, for oversampling, rather than reducing the number of images for larger classes, we opted to duplicate images in smaller classes, ensuring they reached a comparable image count to the larger classes. Following experimentation with both methods, the results indicated that undersampling produced superior outcomes. This is attributed to its ability to mitigate the potential overfitting effects that may arise from oversampling the same set of images.

## Transfer Learning

Due to the small size of the dataset, we performed transfer learning, which is the process of using a pre-trained model, trained on a large and general dataset, to attempt to generalize to new data, in this case, brain microscopy images at the cellular and subcellular level. We tested different pre-trained models including AlexNet (*15*), SqueezeNet (*42*), EfficientNet-b0 (*43*), Inception V1 (GoogLeNet) (*44*), ResNet-18/50 (*45*), and VGG-16/19 (*46*). Inception v1 and ResNet-18 resulted in the best networks. The optimal hyperparameters were found using stochastic gradient descent with momentum, learning rate of 0.001 with a drop factor of 0.1 every 2 epochs, 6 epochs, a mini-batch size of 128, and L2 regularization lambda of 0.0001. We employed early stopping if the loss did not improve after an epoch.

Pre-trained models (such as Inception v1 and ResNet-18) can be fine-tuned or used for feature extraction. Instead of re-training the entire model, fine-tuning involves freezing the initial layers of a model while leaving the top layers, including a newly added classifier layer, unfrozen. When a layer is frozen, the weights are not updated when training on the new data. Earlier layers of a model focus on lower-level features (such as lines and edges), while the top layers extract higher-order features. The top layers are left unfrozen since higher-order features have more relevance when training on new data, while the lower-level features are more common. Feature extraction is the process of using a pre-trained network to extract the features of new data and training a new classifier from those features. Since most pre-trained CNNs perform well when using raw images as the input, training a new classifier on the extracted feature vectors is skipped and training is performed directly by fine-tuning the top layers.

## Model Optimization

When training a model, there are multiple optimizers to choose from. Optimizers are algorithms that are used to update the weights of a neural network to minimize the loss (the penalty for incorrect classifications). In this study, we tested two of the most commonly used optimizers: SGDM (stochastic gradient descent with momentum) and Adam (adaptive movement estimation). SGDM, defined by the equation:

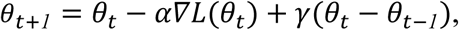

in which: θ = parameters (weights), *t* = iteration number, α = learning rate, *L* = loss function, γ = momentum, updates each weight to minimize the loss function by computing the gradient of the loss function and updates the weight using a random mini-batch. Since SGD (stochastic gradient descent) can oscillate towards the optimum, SGDM implements a momentum term to reduce the oscillation. Unlike SGDM, Adam differs by using learning rates that differ by parameter and can automatically adapt to the loss function being optimized. Adam maintains an average of both the parameter gradients (i) and the square of the parameter gradients (ii):

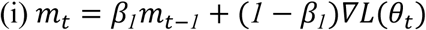

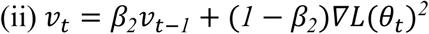

in which: *m_t_* = average of parameter gradients, *v_t_* = average of square parameter gradients, *t* = iteration, β_1_ = gradient decay, β_2_ = square gradient decay, *L* = loss function, θ = parameter. Adam uses the moving averages to update the network parameters:

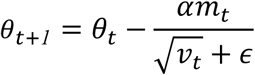

where θ = parameter (weights), *t* = iteration, α = learning rate, *m_t_* = average of parameter gradients, *v_t_* = average of square parameter gradients, ϵ = division by zero constant. Compared to SGDM, Adam can update each parameter based on the gradient’s history. Adam is a faster algorithm compared to SGDM, but it is more susceptible to noise, which may lead to suboptimal performance.

A model must be given the optimal hyperparameters, so that training and testing accuracy converge smoothly, resulting in fitting and avoiding overfitting, therefore resulting in the model learning the key general patterns instead of memorizing (overfitting). Hyperparameters (learning rate, momentum, epochs, mini-batch, etc.) control how the model trains and determine the parameters of the model. In order to optimize the hyperparameters, we used two algorithms: random search and Bayesian optimization. Random search is the process of generating and evaluating random parameters, in this case to minimize loss. Random search is an uninformed algorithm, as it has no knowledge of the previous results. Unlike random search, Bayesian optimization is an informed algorithm. By using the previous results, Bayesian optimization uses a probabilistic model to estimate the next parameters. We used both of these methods to find initial hyperparameters and performed manual tuning to achieve the final hyperparameters.

## Overfitting prevention

Overfitting is a problem in machine learning where a model learns the background noise in the training set and is unable to generalize to the new data in the testing set. The methods we used to combat overfitting are: L2 regularization, dropout, reducing model complexity, and freezing layers. L2 regularization penalizes the model if the weights become too large and encourages the model to have small and a more even distribution in its weights. L2 regularization is included in the hyperparameters that can be tuned to minimize the loss (difference between the predicted value by the model with the original value given in the dataset) thereby the networks generalize better to new data (*47*). Dropout is another regularization method that can be implemented to different layers within a CNN. By implementing dropout, a different node within a CNN will randomly be disabled (or “dropped”) with probability *p*. Dropout helps combat overfitting by forcing the model to learn more robust features, which can improve generalization. If a model picks up the background noise and features of the data, it can be a sign that the model is too complex for the task at hand, and removing layers to create a simpler model can help with overfitting. Freezing the layers can also help reduce overfitting by limiting the number of parameters that are updated when training, also reducing the model complexity.

### Analysis, Statistics, and Visualization of Results

## Evaluation of Model Performance: K-Fold Cross Validation

We used several independent methods to estimate the accuracy of the model and analyze findings. Cross validation is a general method used to determine the accuracy of a trained machine learning model on unseen data. In this study, we used k-fold cross validation, which provides a more reliable performance evaluation compared to a train-test split (*48*). For k-fold cross validation, we divided the original dataset into *k* equal-sized partitions, or “folds.” The model was trained *k* times, each time using *k*-1 folds as the training data, and the remaining fold as the testing data. This ensured that each fold was used once for testing. In our design, each fold was a case, and different cases were rotated between the training and testing sets. The performance of each model was evaluated using metrics such as accuracy or other precision and recall metrics. By averaging the results from each model, an estimation of the overall performance of the model was obtained. By using k-fold cross validation, the model was trained and evaluated on different subsets of the data, facilitating assessment of the generalizability of the model.

## Evaluation of Model Performance: Classification Confusion Matrices

We additionally used confusion matrices to evaluate the accuracy of the network after training. In confusion matrices, the rows display the true classes, while the columns display the predicted classes, or what the network predicts which class the images belong to. For multiclass confusion matrices, the on-diagonal displays the correctly classified images, also known as true positives. The values in a column for a specific class that is not the true positive reflects the false positives for that class, the values in the row that is not the true positive reflects the false negatives for that class, and all other values not in the corresponding row and column for that class reflect the true negatives. We additionally combined the 4 original classes (ASD SWM, ASD DWM, CTR SWM, and CTR DWM) into ASD vs CTR and DWM vs SWM to analyze the network performance for classification of pathology and white matter region/depth, respectively.

## Evaluation of Model Performance: Estimation of Precision and Recall Metrics

To take into account false positives and negatives in our accuracy measures we estimated two additional evaluation metrics, precision and recall, which in image classification tasks provide in depth analysis of the performance of a machine learning model (*49*). For multiclass classification, precision, also known as positive predictive value, is defined as the fraction of correct predictions (true positives) over all predictions (true positives and false positives) for a single class. A higher precision signifies a low prevalence of false positives, and when the network predicts a certain class, it is likely to be correctly classified. Recall, also known as sensitivity or true positive rate, is the fraction of true positives over all actual positives (true positives and false negatives). A higher recall score signifies a low prevalence of false negatives (misses), and that the network can accurately identify instances of true positives. By calculating the harmonic mean of both precision and recall, the F1 score is obtained. Compared to standard accuracy, F1 score provides a more balanced metric, taking into account false positives and negatives.

## Evaluation of Model Performance: Receiver Operating Characteristic Curves

A second performance evaluation metric that we implemented is the receiver operating characteristic curve (ROC curve), which shows the performance of a neural network at different classification thresholds/criteria. An ROC curve plots the true positive rate (TPR) vs false positive rate (FPR), and by altering the classification threshold, the true positive rate and false positive rate can increase or decrease, depending on the classification threshold. The best operating point can be chosen based on the desired true positive and false positive rates. Another aspect of a ROC curve is the area under the curve (AUC). The AUC provides an aggregate measure of performance across all possible classification thresholds and represents the network’s ability to differentiate between true and false positives. The closer the AUC is to 1.0, the better the model’s ability to separate classes from each other.

## Image Sensitivity Maps

To further analyze and visualize the results we generated image sensitivity maps using the following two approaches: Occlusion sensitivity maps, also known as saliency maps, provide insights into how the network creates its predictions by identifying the salient regions of the image that positively and negatively contribute to the network’s classification. Occlusion sensitivity maps operate by sliding an occluding mask across the input data, resulting in a change in the classification score at each mask location. This process enables the identification of specific areas within the input data that exert the most influence on the classification score. In addition to experimenting with the default size and stride for the occluding mask (with a mask size of 20% of the input size and a stride of 10% of the input size), we explored various combinations of stride and size to determine the optimal patch for highlighting axonal details.

Gradient class activation mapping (Grad-CAM) is a second approach utilized to provide insight into the regions of interest for network predictions. Grad-CAM, which is based on the CAM technique, determines the importance of each neuron in a network prediction by considering the gradients of each target classification flowing through the deep network. This results in a localization map highlighting the regions of interest for the desired class. Compared to occlusion sensitivity, Grad-CAM provides a more general overview of the features that contribute to the class, while occlusion sensitivity is more specific to what specific areas the network identified to make its prediction. Grad-CAM can be used in similar ways as occlusion sensitivity, but these two image sensitivity maps are best used complementary to find generalizations and specializations, respectively. Using both approaches, we analyzed the overlaps and lack of overlaps in the regions of interest of the model vs. significant features of ACC white matter from prior neuroanatomical studies (*1–3*). Our aim was to use image sensitivity maps to decipher the aspects of network focus, so that we can then a) identify novel features and metrics that have not been studied extensively before or have not reached significance in prior comparisons and, b) identify features that were not emphasized by the network but featured prominently in prior neuroanatomical studies.

When graphing image sensitivity images, we also included the probability predicted by the network for that class to show the level of certainty the network had for the classification of a certain image. In addition to producing heat maps for the specific class (e.g. ASD DWM, ASD SWM, etc.), we also included the combined grad-cam for each pathology, such as combining ASD DWM and ASD SWM for a complete ASD representation. Compared to other saliency map techniques such as occlusion sensitivity, Grad-CAM provides a more global and broader outlining view of regions that contribute most towards classification, while other position-based saliency maps such as occlusion sensitivity provide a more local view (Supplementary Figure 1).

## Generation of DeepDream Images

To better visualize the data and highlight patterns that were characteristic for each group and played a key role for correct classifications we generated DeepDream images. DeepDream, a computer vision algorithm created by Google in 2015, enhances the features found in images and creates visualizations of the features extracted by a neural network for a desired classification (*50*). After a network is trained, the DeepDream algorithm alters an image consisting of random noise to maximize “loss”, or the sum of activations for a chosen layer of a neural network to excite the layer for a desired class. Earlier layers may produce DeepDream images that consist of simpler patterns such as edges or colors, while later layers are more detailed and include sophisticated features. When training a network, loss is typically minimized through gradient descent, but the DeepDream algorithm maximizes loss through gradient ascent for a specified class. By generating DeepDream images from the training data, the enhanced features can be noted for each class, cross validated with image sensitivity maps, and can be used as guidelines for future classifications.

## Principal Component Analysis

To extract and analyze relevant image features and metrics for each classification of the CNN we applied dimension reduction techniques, typically used in conjunction with machine learning approaches (*21, 51*). For each convolution layer in a neural network, an activation function is used to convert the input values into an output. In terms of image classification, the output is typically a value that corresponds to a specific class (*15, 52*). The fully connected layer, which receives inputs from all its previous layers, performs a weighted summation on the class-specific activations, and feeds the values into a SoftMax activation function that normalizes the values into probabilities to determine the final output class. By utilizing the layer activations from the fully connected layer, each input, represented by a layer activation vector, can be analyzed for its similarities and uniqueness for each class. For multiclass classification tasks, the activation vectors can contain N dimensions, but the relevant information is contained in a smaller M dimension. To extract the relevant M dimensions, dimension reduction techniques are applied, and in this study, we specifically used principal component analysis (PCA), to better analyze and visualize the data. PCA is an optimal dimension reduction technique that captures the maximum variance in the data through singular value decomposition (SVD). Through this process, features that provide no information or negligible information are less represented, and the best low-dimensional approximation of the data is constructed. In order to perform PCA, the original data matrix was normalized, the covariance matrix was computed and then decomposed into its eigenvalues. Given an *m* x *n* matrix **M**, the standardized matrix **Z** was computed as:

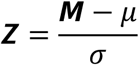

where μ is the mean and σ is the standard deviation. The covariance matrix **C** of **Z** is given by:

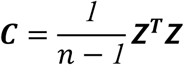

where n is the number of observations in the data. Eigenvalue decomposition of **C** yields:

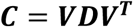

where **V** is the matrix of eigenvectors and **D** is the diagonal matrix of eigenvalues. The eigenvectors were ordered by their corresponding eigenvalue in descending order, which can be used to determine the importance of each principal component. After performing eigenvalue decomposition, the loadings were also calculated, which indicate how much each variable contributes to each principal component and helps understand the meaning of the principal components in terms of the original variables. After eigenvalue decomposition, the loadings matrix **L** was calculated by:

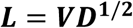

where **D^1/2^** is the diagonal matrix, whose diagonal elements are the square roots of the eigenvalues. Each column of the loadings matrix **L** corresponds to a principal component, and each row corresponds to one of the original variables. The magnitude of a loading indicates the importance of the corresponding variable to the principal component, and the sign of a loading indicates the direction of the relationship between the variable and the principal component.

Compared to other dimension reduction techniques such as MDS, PCA does not use the Euclidean distance of activations, but the value of the activations from the layers themselves. PCA increases the interpretability of higher dimensional data while preserving the key features of the data as specified by the singular values obtained from SVD.

### Code/Software

All codes and analyses are deposited and available in https://github.com/kkuang0/ASD-ACC-Transfer-Net(public access).

## Results

### DNNs Perform Well with Pathology Classification and Single Out ASD SWM

We partitioned the dataset into three subsets: a training set for network training, a validation set for periodic progress assessment during training, and a testing set to evaluate the final network’s accuracy. This data-splitting procedure was repeated multiple times, each time using different case combinations for training, validation, or testing. Table 2 presents the training results for all combinations, ranked from highest to lowest pathology classification accuracy.

The highest-performing network for pathology distinction using EM images achieved an accuracy of 52.65% across four classes (Figure 2A.i). Notably, focusing solely on pathology classification resulted in significantly improved performance, with the highest accuracy reaching 90.88% for EM images (Figure 2A.ii). For Tol-Blue optical microscopy images, the best-performing network for pathology distinction achieved an accuracy of 67.52% across four classes (Figure 2B.i) and 97.68% for pathology (Figure 2B.ii). Conversely, classification accuracy for DWM vs SWM was lower, with 57.21% for EM images (Figure 2A.iii) and 69.13% for Tol-Blue images (Figure 2B.iii). Therefore, the major source for the low 4 class accuracy stems from the DWM vs SWM confusion.

**Figure 2:**
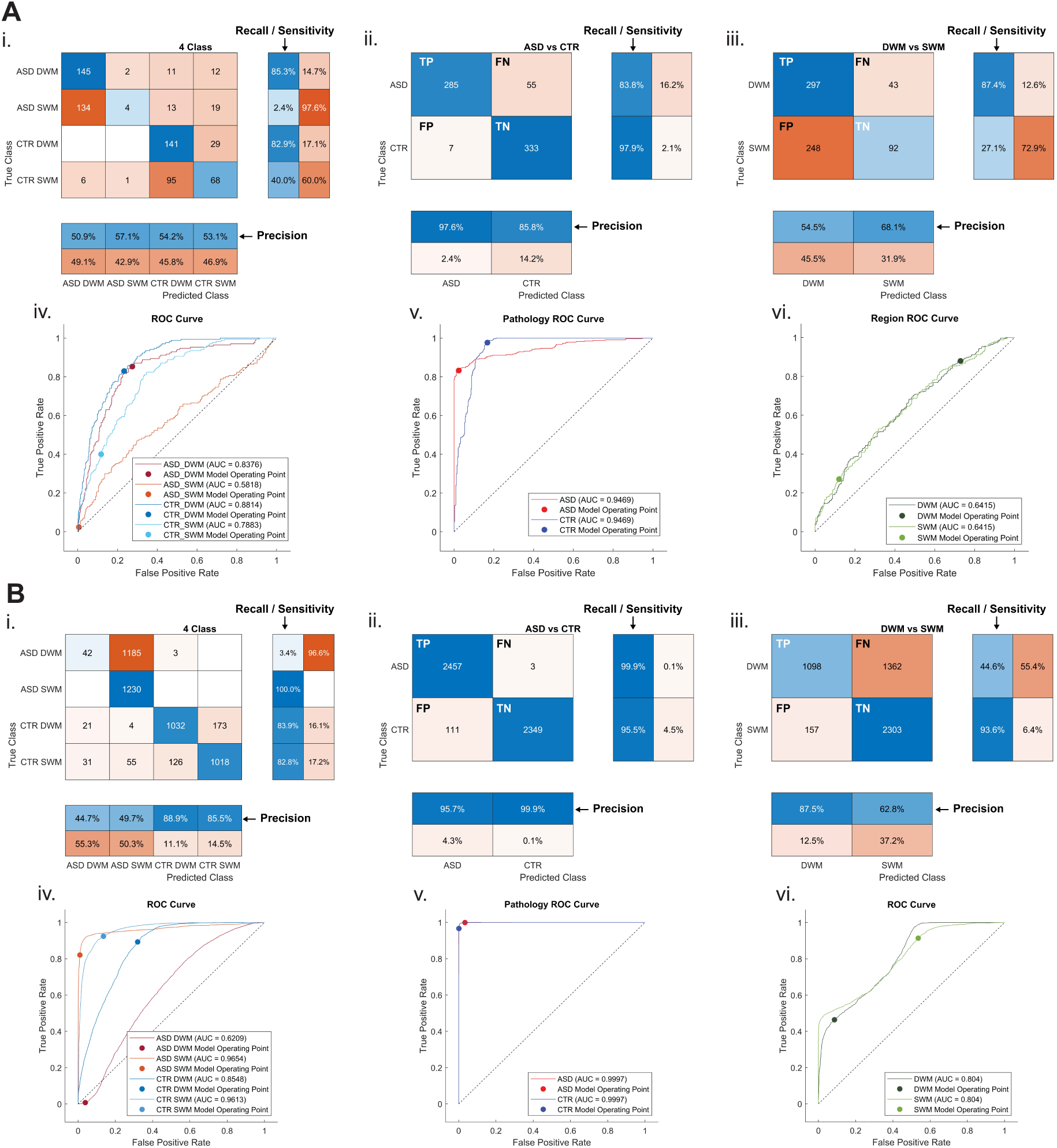
Confusion matrices and ROC curves for model classification. **A.** Classification of EM images. **B.** Classification of Tol-Blue microscopy images. For both panels A and B: **i.** 4 class confusion matrix of pathology and white matter region, **ii.** white matter depth independent pathology matrix, **iii.** pathology independent white matter depth matrix, **iv.** 4-class ROC, v. white matter depth independent ROC, **vi.** pathology independent white matter depth ROC. Confusion matrices show the true classes compared to the predicted classes. For 4-class confusion matrices (A.i, B.i), the column summaries correspond to precision, and the row summaries correspond to recall for the corresponding classes. For 2-class confusion matrices (A.ii, A.iii, B.ii, B.iii), the confusion matrix cells (from left to right, top to bottom) correspond to true positive (TP), false negative (FN), and false positive (FP), and true negative (TN) (with respect to ASD for panels marked ii., and to DWM for panels marked iii.). An ROC curve plots the true positive rate (TPR) against the false positive rate (FPR) and shows the performance of a classification model at all classification thresholds. The model operating points represent the threshold that minimizes the FPR and maximizes the TPR.

For EM images, the classes that had the highest accuracies were ASD DWM and CTR DWM, 88.8% and 96.5% respectively (Figure 2A.i) Conversely, the class with the most misclassifications was ASD SWM with most of them being misclassified as ASD DWM, with an accuracy of only 14.7%. This aligns with the characteristics of patients with ASD, whose SWM is typically the most variable (*1, 2, 4*).

For Tol-Blue images, the network perfectly classified ASD SWM, but also classified 96.57% of ASD DWM slices as ASD SWM (Figure 2B.i). Of all slices classified as ASD, 98.29% were classified as ASD SWM. This suggests that the network mainly learned the features of ASD SWM and generalized those features as ASD. The network could distinguish between different CTR white matter depths with similar accuracies: 83.9% for DWM and 82.8% for SWM.

### Dimension Reduction Graphs Further Support the Classification Results

To visualize the dataset in three-dimensional space and observe the variance among the data points, we extracted the feature vectors from the last fully connected layer (see Methodology) to generate 3D PCA graphs from the best-performing network for EM and Tol-Blue images (Figure 3).

**Figure 3:**
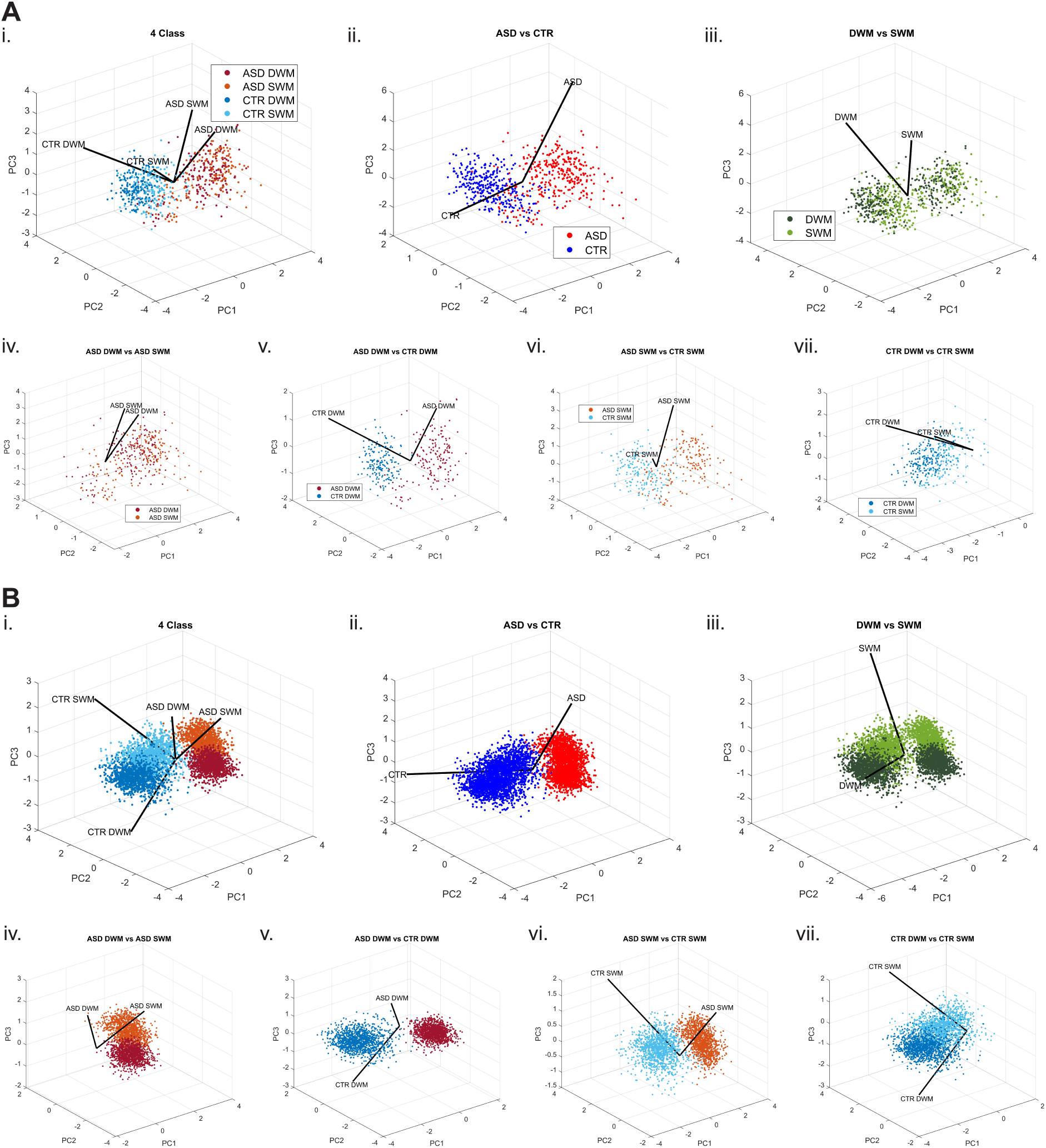
PCA biplots of EM and Tol-Blue microscopy images. **A.** EM analysis. **B.** Tol-Blue analysis. **i.** 4-class, **ii.** ASD vs CTR, **iii.** DWM vs SWM, **iv.** ASD DWM vs ASD SWM, **v.** ASD DWM vs CTR DWM, **vi.** ASD SWM vs CTR SWM, **vii.** CTR DWM vs CTR SWM. The axes correspond respectively to the first, second, and third principal component. PCA loadings describe how much each variable contributes to a particular principal component and the variance explained by that principal component. Plots for pooled categories, in ii. and iii., highlight that the higher accuracy of the classification for pathology, as opposed to white matter depth, showing that the ASD versus CTR data points were significantly more dispersed and distinct, whereas the white matter depth data points were more interspersed and less distinguishable.

For EM images, consistent with the confusion matrices, the PCA plots clearly distinguished between the ASD data points and the CTR data points (Figure 3A.i, Figure 3A.ii.), when viewing the first three principal components. On the other hand, the data points for DWM vs SWM were more mixed in with one another, indicating a higher confusion for white matter layer classification (Figure 3A.iii.) The loading vectors for the classes also showed a similar trend as ASD and CTR vectors appeared further apart and more distinct (angle > 90 deg, Figure 3A.ii.) compared to the SWM and DWM vectors (angle < 90 deg, Figure 3A.iii.), suggesting differences in the relationship between each variable and the principal component. That is because the magnitude of a loading indicates the importance of the corresponding variable to the principal component, and the sign of a loading indicates the direction of the relationship between the variable and the principal component.

For Tol-Blue images, the PCA plots showed similar results as the EM images, with a clear separation between ASD and CTR, but less distinction between the white matter depths (Figure 3B.i, Figure 3B.ii., and Figure 3B.iii.). When comparing the loadings for pathology (Figure 3B.ii.) and white matter depth (Figure 3B.iii.), the ASD and CTR loadings had an angle of 99.823 deg (1.742 rad), while the DWM and SWM loadings had an angle of 94.722 deg (1.653 rad), which is consistent with better white matter depth separation in Tol-Blue compared to EM images.

Pairwise plotting from 2 of 4 total classes also reveals similar results, with the most confusion coming from the DWM vs SWM pairs (ASD DWM vs ASD SWM and CTR DWM vs CTR SWM) (Figure 3A.iv, Figure 3A.v., Figure 3B.iv, Figure 3B.v.). Figures 3A-3B, vi and vii clearly show the distinction between the ASD and CTR groups, whether it’s ASD DWM vs CTR DWM, or ASD SWM vs CTR SWM as the data points in each respective group are spread apart from left to right.

### Principal Component Analysis Emphasizes Variance and Correlations

To understand the importance of each principal component, we also estimated the percentage of total variance explained by each component. In EM images, the first principal component captured 70.01% of the explained variance, while the second principal component added in another 14.78% (Figure 4A.ii). Similarly, in the Tol-Blue slices, we observed that 70.1% of the explained variance was captured by the first principal component and an additional 25.3% by the second principal component (Figure 4B.ii), which indicates that the variability of the data was successfully captured by the first two principal components. We chose the third principal component to be the cutoff to maximally contain as much explained variance (94.11% and 98.8% for EM and Tol-Blue, respectively), while still reducing the dimensions to facilitate interpretation.

**Figure 4:**
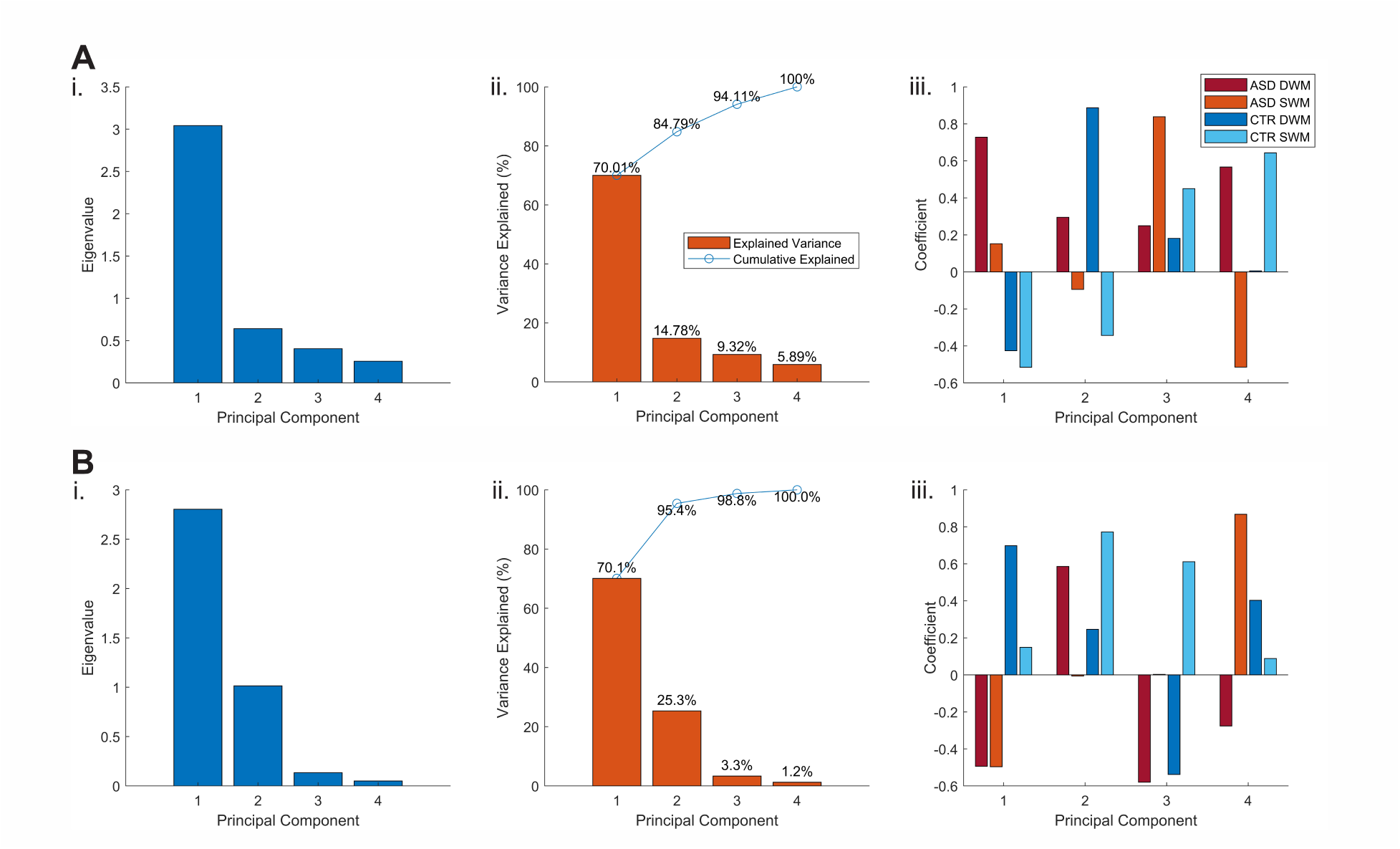
Scree and loading plots after PCA of EM (A) and Tol-Blue (B) microscopy images. **i.** Eigenvalues of each principal component, **ii.** explained variance by each principal component and cumulative explained variance by all principal components, **iii.** coefficients of each class contributing to each principal component.

The coefficients for each principal component, known as loadings, also revealed crucial details of how the network viewed the data. Each loading vector represents how much each variable contributes to a principal component, while the direction and magnitude of each vector indicate correlations and magnitudes between each class and principal component. The different coefficients of each loading indicate how each class contributes to the component. In the first principal component loading, ASD classes and CTR classes had different signs (Figure 4A.iii, 4B.iii). For EM images, both ASD DWM and ASD SWM had positive coefficients: 0.7279 and 0.1523, while their corresponding CTR classes were negative: -0.4256 and -0.5155 (Figure 4A.iii). We saw the inverse for Tol-Blue slices, where both ASD classes had negative coefficients, and the CTR classes had positive coefficients (Figure 4B.iii). This indicates that the first loading feature corresponded to the separation between ASD and CTR. This also explains why there is such a clear distinction between the ASD and CTR classes in the scatter plots (Figure 3), as the variance between ASD and CTR is mainly captured. The distinction between DWM and SWM regions was also reflected in the loadings. In the second principal component loadings for EM images (Figure 4A.iii), the DWM regions were positive (0.2951 and 0.8865), while the SWM regions had negative values (−0.4257 and -0.5155). The same was true for Tol-Blue images for its third principal component (Figure 4B.iii), as Tol-Blue’s loading had negative coefficients for the DWM regions, and positive coefficients for the SWM regions, indicating that this feature was responsible for the distinction between white matter regions. In the third principal component loading for EM, all the coefficients were positive, with a tendency for the loadings of DWM regions to be closer to zero (0.2490 and 0.1810), while the loadings from SWM were larger (0.8384 and 0.4498); the sizable difference between ASD SWM’s loading compared to other classes points out the confusion about ASD SWM. This is consistent with the confusion matrix as the network clearly struggled to identify ASD SWM correctly and mostly misclassified it as ASD DWM (Figure 2A.i). There was a reverse confusion as well in Tol-Blue images, where the network consistently confused ASD DWM as ASD SWM (Figure 2B.i). This can be explained by the second principal component, as the coefficient for ASD SWM is significantly smaller compared to the other classes, indicating that the second principal component mainly differentiated the other classes from ASD SWM (Figure 4B.iii). We also found that the explained variance for the principal components responsible for DWM and SWM separation only covered 14.78% for EM, and 3.3% for Tol-Blue (Figure 4A.ii, 4B.ii). This can explain why the separation between DWM and SWM was less clear compared to pathology classification.

### Heat Maps Reveal Crucial Areas for Classification

We used heat maps produced by Occlusion sensitivity and Grad-CAM approaches to analyze and highlight the overlaps or lack thereof in the regions of interest that weighed most for the model classification output and compared with significant features and pathology of ACC white matter from prior neuroanatomical studies (*1–3*). Grad-CAM overall provided a more global and broader outlining view of regions that contributed most towards classification that was easier to interpret, whereas occlusion sensitivity provided a more local view that was in many cases too narrow and difficult to interpret. Grad-CAM-generated saliency maps for EM (Figure 5) and Tol-Blue images (Figure 6) consistently highlighted the saliency of axon density, size, and trajectory variability as key features for correct classification. Importantly, hotspots that included cells (mainly nuclei of glia) and relatively uniform axon regions were typically salient in incorrectly classified images (Supplementary Figure 2).

**Figure 5:**
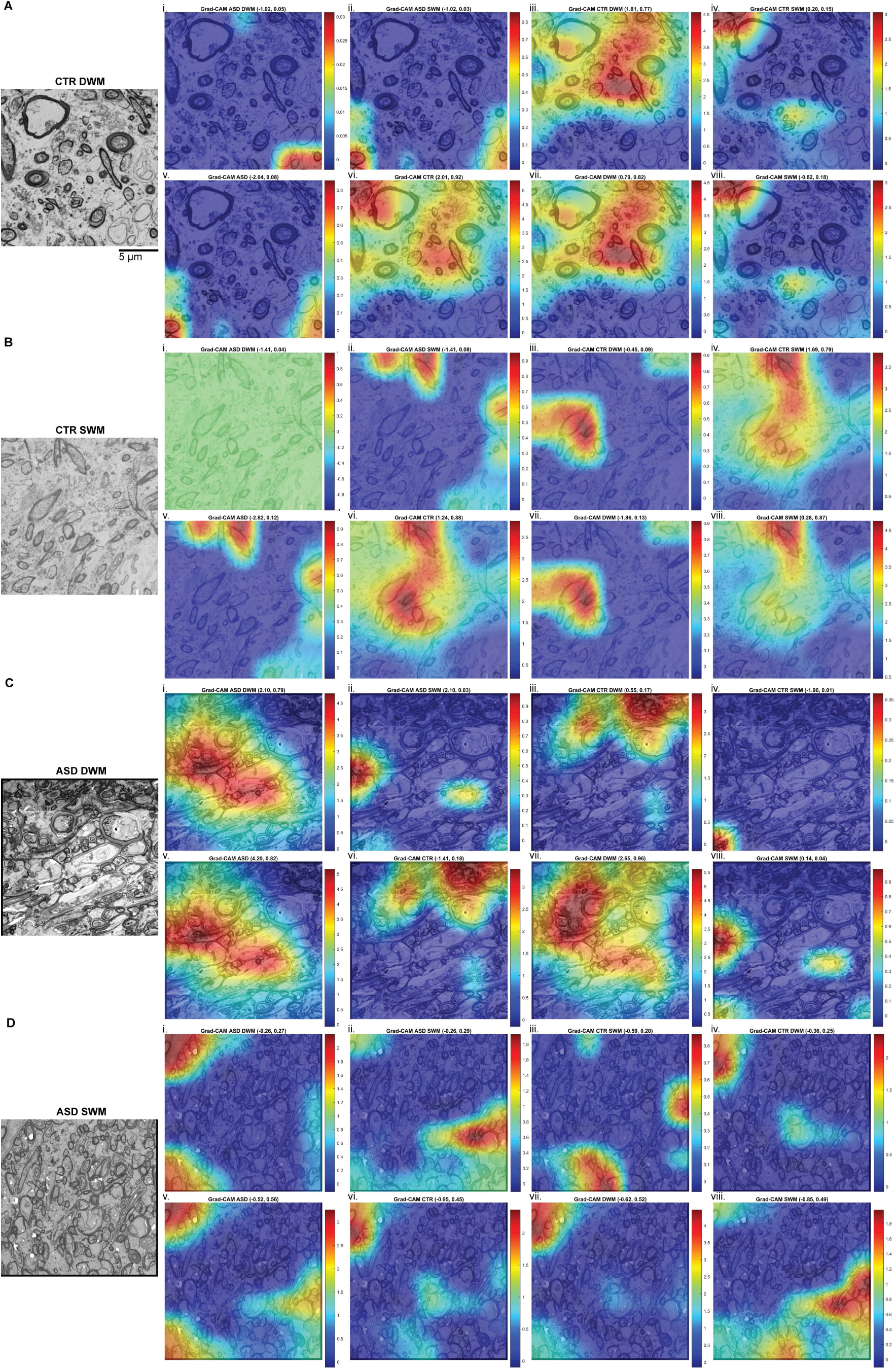
Image sensitivity maps of correctly classified EM images. **A-D**, Representative examples of EM images and overlaid heat maps from each of the 4 classes: CTR DWM (A), CTR SWM (B), ASD DWM (C), ASD SWM (D). For each class we used Grad-CAM to generate 8 heatmaps, representing hotspots and their relative weights (probability) for correct and incorrect class assignment: **i.** ASD DWM, **ii.** ASD SWM, **iii.** CTR DWM, **iv.** CTR SWM, **v.** ASD, **vi.** CTR, **vii.** DWM, **viii.** SWM. The title of each heatmap shows the sum of model neuron activity from the fully connected layer, as well as the SoftMax probability. Heatmaps of the correct corresponding class highlight areas of the image that contributed most towards the overall classification. Heatmaps of the other classes provide a comparative visualization of how different features were weighted. This aids in identifying specific patterns and features that distinguish each individual class, enhancing the interpretability and transparency of the network’s performance.

**Figure 6:**
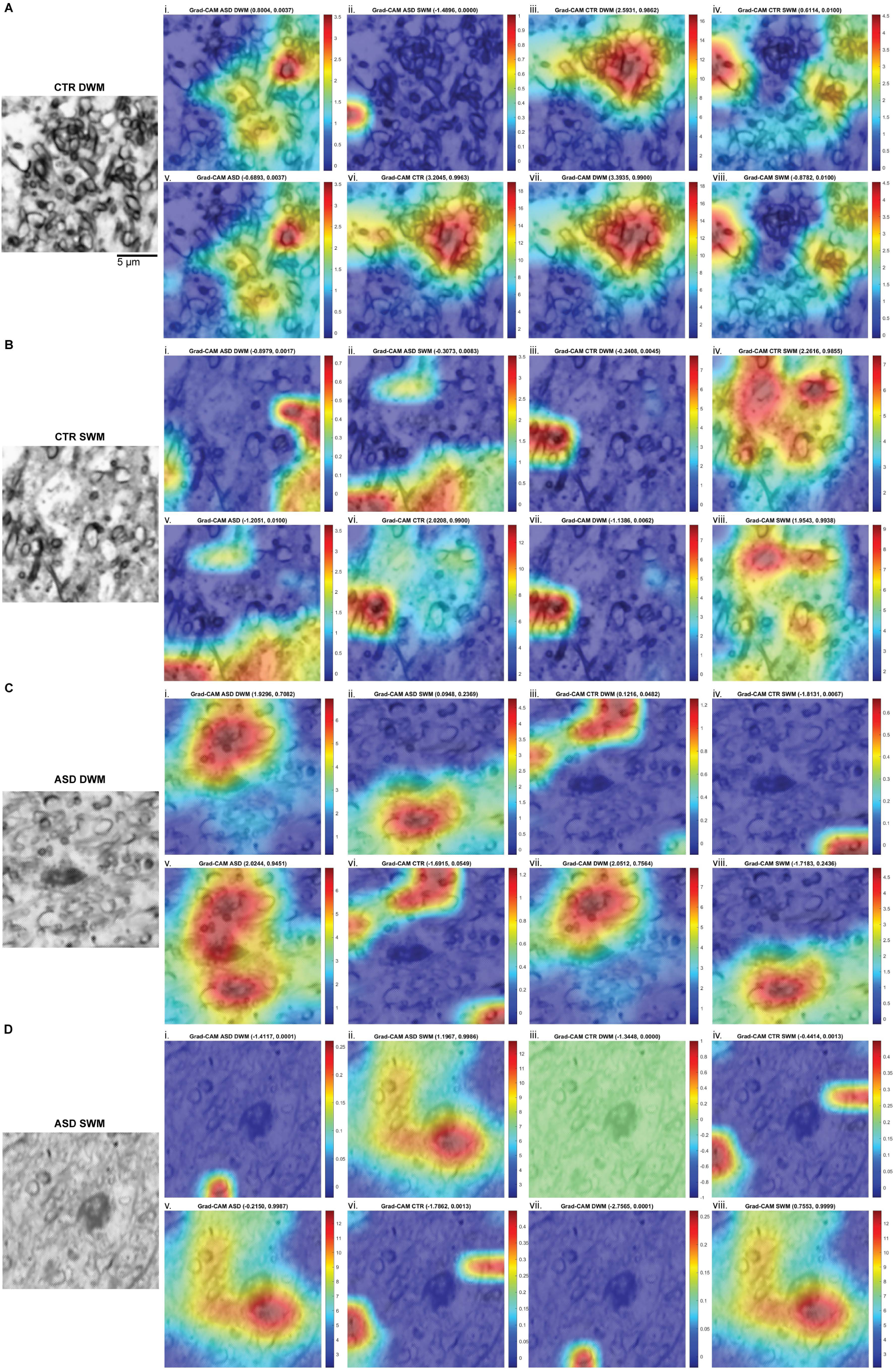
Image sensitivity maps of correctly classified Tol-Blue microscopy images. **A-D**, Representative examples of Tol-Blue images and overlaid heat maps from each of the 4 classes: CTR DWM (A), CTR SWM (B), ASD DWM (C), ASD SWM (D). For each class we used Grad-CAM to generate 8 heatmaps, representing hotspots and their relative weights (probability) for correct and incorrect class assignment: **i.** ASD DWM, **ii.** ASD SWM, **iii.** CTR DWM, **iv.** CTR SWM, **v.** ASD, **vi.** CTR, **vii.** DWM, **viii.** SWM. The title of each heatmap shows the sum of model neuron activity from the fully connected layer, as well as the SoftMax probability.

Heatmaps of the correct corresponding class highlight areas of the image that contributed most towards the overall classification. Heatmaps of the other classes provide a comparative visualization of how different features were weighted. This aids in identifying specific patterns and features that distinguish each individual class, enhancing the interpretability and transparency of the network’s performance.

### DeepDream Images Generalize the Extracted Features from Each Class

DeepDream is an iterative process that alters an image of random noise to maximally increase the confidence level for the specified output class (*50*). For both EM and Tol-Blue images, the DeepDream images for DWM regions of both ASD and CTR (Figure 7A.i., iii., and 7B.i., iii.) showed distinct axon-like bundles and clustering, which is a key defining feature of DWM regions. On the other hand, the SWM DeepDream images appeared more convoluted, indicating that SWM regions contained a variety of axonal profiles with heterogeneous trajectories, including cross-oriented and perpendicular axons (Figure 7A.ii., iv., and 7B.ii., iv.). When comparing ASD and CTR images, the CTR images showed higher axonal orientational consistency in both DWM and SWM regions compared to the respective ASD regions.

**Figure 7:**
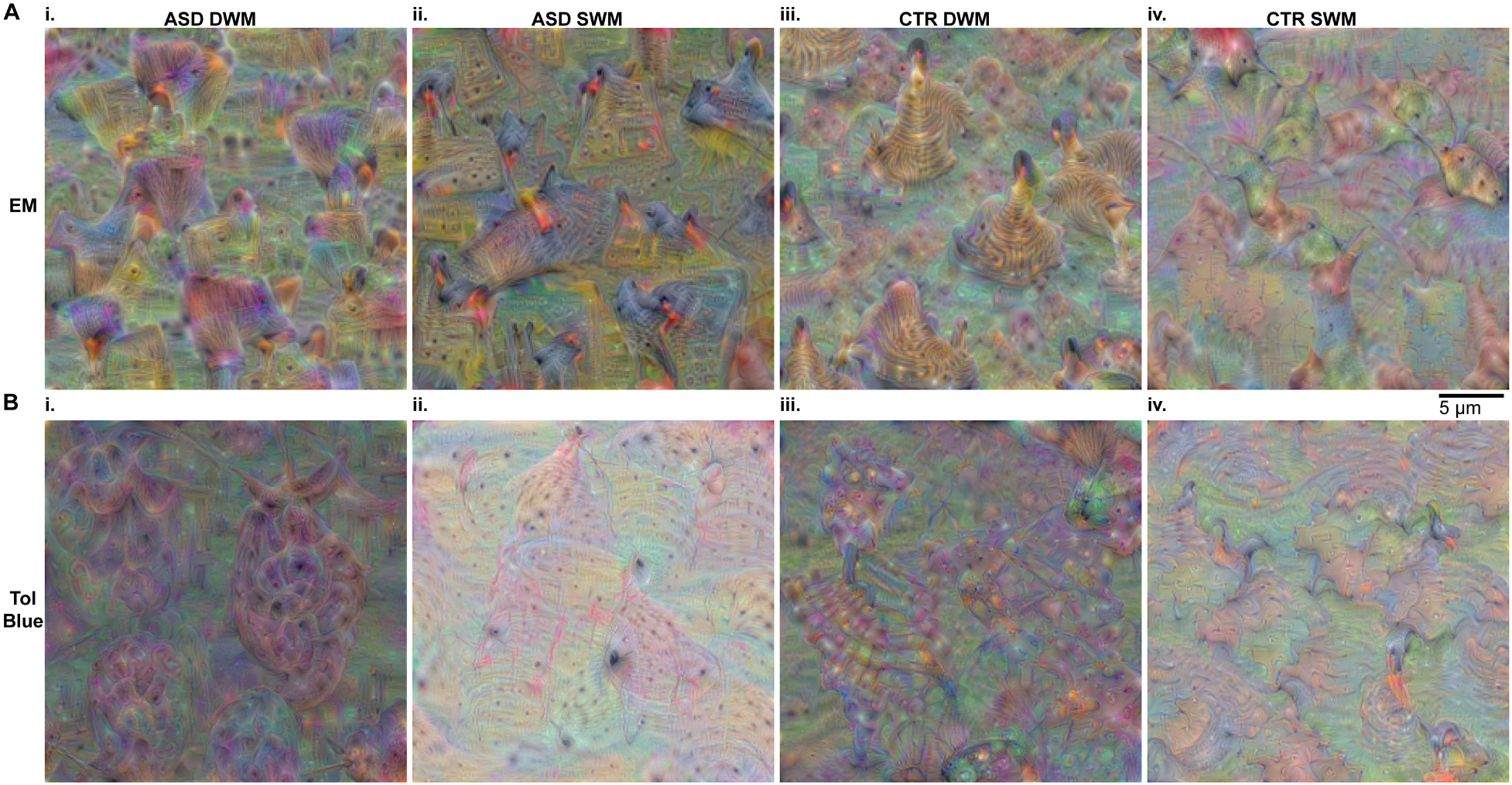
DeepDream images can generalize the salient extracted features from each class. **A.** EM and **B.** Tol-Blue DeepDream images constructed from the networks for each class: i. ASD DWM, ii. ASD SWM, iii. CTR DWM, iv. CTR SWM. EM DeepDream images showed the network’s maximally amplified activations for EM features, highlighting the textures the network extracted for each class. Tol-Blue DeepDream images similarly showed the network’s maximally amplified activations for Tol-Blue features, highlighting more of the structures as well as textures the network extracted for each class. In the DWM sections for both EM and Tol-Blue, the DeepDream images created distinct axon bundles, while the SWM images contained sparse, scattered axons of different orientations. These DeepDream images offer insights into the internal representations and feature hierarchies learned by the network for each class for accurate classification.

Specifically, the CTR DWM image showed more organized distribution patterns of axons, while the ASD DWM image contained higher randomness of axon distribution. When comparing the ASD DeepDream images for EM and Tol-Blue, we noticed that the EM DeepDream images were characterized by a checkered (square-like) pattern, while the Tol-Blue images were characterized by wavy patterns. This could reflect patterns observed in the EM and Tol-Blue Grad-CAMs for ASD SWM (see Figures 5, 6), where the axons in the areas highlighted by the heatmaps in EM formed a more rectangular structure, while the Tol-Blue Grad-CAMs mostly highlighted parallel axons, creating the distinct bundles.

## Discussion

High-resolution quantitative study of features of individual white matter axons in multiple cortical pathways can provide key insights about pathway features, integrity, signal transduction, and neural communication in the neurotypical brain and mechanisms of disruptions in ASD. However, such analyses require extensive study by experienced anatomists and semi-automated tracing of individual axons. They are thus labor-intensive and time-consuming, rendering these approaches not optimal for large-scale studies that can help identify core ASD network status and likely mechanisms of disruption in communication. A handful of studies have successfully undertaken this task (*4–6, 8, 53, 54*) however, there is a critical need for more detailed studies that will survey relevant cortical and subcortical pathways in neurotypical healthy controls and individuals with ASD. To address this need we developed a multidisciplinary approach that combines sophisticated high-resolution quantitative microscopy and histopathology with advanced methodologies of machine learning and artificial intelligence (AI). We used our large dataset of light and electron microscopy images of white matter axons from short- and long-range cortical pathways below the anterior cingulate cortices in control and ASD groups of adults, from studies published since 2010 (*4, 53, 54*), to develop and optimize machine learning algorithms. Our results showed that by customizing and optimizing deep neural networks (DNNs) through transfer learning, we can determine if microscopic images were obtained from *post-mortem* brains of ASD or typically developed individuals, with high confidence (98%). Moreover, we were able to extract image sensitivity maps, to highlight key features in the images that contributed to the accuracy and errors of DNN classification, and evaluate the extend of these contributions, revealing novel insights about mechanisms of ASD pathology.

Overall, the final trained networks demonstrated distinction across four classes (ASD DWM, ASD SWM, CTR DWM, and CTR-DWM) at multiple scales, spanning cellular (Tol-Blue) and subcellular (EM) levels. Loading values from principal components revealed crucial details about how the networks learned. The first principal component for both types of images detected the difference between ASD and CTR, which explains the high accuracy of our networks in classifying pathology. The second principal component was primarily focused on white matter region classification. This was reflected in the network’s moderately successful performance in classifying white matter regions. Classification across two categories (ASD vs CTR or SWM vs DWM), after pooling across a factor (e.g., DWM and SWM) to assess predicted image classes (e.g., ASD vs. CTR) against the true class of each image, bumped up accuracy significantly. Importantly, when the classification results were pooled together across pathology (ASD vs CTR) and depth (deep vs superficial white matter), the networks consistently exhibited a better performance in classifying pathology compared to white matter regions (Figure 3 and Table 2). Notably, the accuracies for ASD detection in both EM and Tol-Blue images were very high, but for different white matter regions, when pooled across different white matter depths. Specifically, in EM images, the network predominantly identified all ASD slices as ASD DWM, whereas the Tol-Blue network primarily classified ASD slices as ASD SWM. This shift in bias in different networks suggests confusion, particularly regarding axons across white matter depths in ASD brains. The difficulty in distinguishing white matter depth led the networks to either predominantly learn traits from ASD DWM and classify these as ASD for EM images or, for Tol-Blue, learn traits from ASD SWM and generalize those features as ASD. This confusion aligns with known anatomical differences, as the superficial and in some cases the deep white matter in patients with ASD tend to significantly deviate from the superficial and deep white matter pattern of neurotypical brains (*1, 2, 4, 6, 7, 13, 19, 55–65*). Supporting this observation, the loading in the third principal component for ASD SWM, in EM images, was the highest, showing the most dispersion compared to other classes, and indicating that the network identified more variability within ASD SWM (Figure 2). On the other hand, in the second principal component for Tol-Blue images, the ASD SWM coefficient was the lowest, and an outlier, showing again separation of ASD SWM from the other classes (Figure 2). Both trends, suggest that white matter depth classification accuracy is more challenging in ASD, compared to controls, and establish the central role of SWM changes under ACC in ASD, providing novel insights about a key locus of heterogeneity and pathology in the white matter below ACC.

Importantly, both correct and incorrect classification of images can provide useful insights for the organization of the white matter below ACC in neurotypical individuals and in ASD. The transition between SWM and DWM is gradual and is evident in the coronal sections used here, by the gradual change in the orientation, trajectory variability, and density of axons, as described in previous studies (*1, 2, 4*). Our findings showed that in the ASD group, in particular, the distinction between SWM and DWM was difficult, a novel insight which suggests blending of the white matter regions, due to widespread changes in axon features. Once the DNN learned these features in the SWM or DWM they were then generalizable to all white matter and necessary for ASD classification. Generated image sensitivity maps highlighted specific attributes of images that contributed to DNN classification, providing an opportunity to compare the focus of DNN on each image with key loci and attributes important for neuroanatomical image analyses. This comparison pinpointed areas of overlap, misses, or novel attributes, converging on the combination of three key features for pathology classification: axon density, heterogeneity of axon sizes, and axon profile shape variability, all of which appeared to be consistently higher in ASD below ACC.

### Limitations, Challenges, and Future Directions

It is well established that axons in the white matter below ACC change in ASD, therefore it is reassuring and not surprising that this machine learning approach could classify the images with high accuracy, after optimization and training of the DNN. A DNN is an artificial neural network with multiple layers between its input and output layers and has simulated neurons, synapses, weights, thresholds, and signal functions. Nonlinear feedforward DNNs with three or more layers are considered “universal approximators”, i.e., with the proper architecture and dataset, they can learn to detect and recognize patterns with high accuracy (*66*). However, the success of a DNN can be significantly constrained by limited training data and limited computational power to perform the training. Moreover, DNNs perform pre-designed computations, such as convolutions (hence the interchangeable term convolutional neural network, CNN) that can yield valuable properties such as image position invariance to the classification power, but the presence of such pre-designed computations, beside their merits, can limit the DNN domain of universality (*67*). We used multiple methods including DNN transfer learning and augmentation through additional pre-processing of images, fine tuning of the number of epochs, and iterations to maximize the accuracy of performance for the images of interest, in line with previous studies (*68–73*). Additionally, we evaluated network performance using dimension reduction techniques, to visualize outliers for further inspection, and using image sensitivity maps, to identify regions and patterns causing DNN confusion or to highlight novel attributes that can be examined. Our findings highlight that even when multiple attributes were combined, to form mixed labels (such as ASD-DWM) for DNN supervised training, the performance of the network for individual attributes (such as ASD vs. CTR) could be redeemed, paving the way for iterations using a multitude of combined labels, as needed. Our exhaustive training and testing regime with different combinations of non-overlapping cases (Table 2), also revealed that with small datasets, in this case limited number of cases and images, it was best to train separate models for optimal classification of pathology only, or white matter depth only, or combinations of these. Compilation of larger datasets with increased number of cases and images that can be used for future training and validation can further maximize performance and increase generalizability.

Other recent studies that use machine learning have similarly focused on brain network connectivity, either functional, based on resting-state functional magnetic resonance imaging (rs-fMRI) and EEG (*18, 51*), or structural, based on MRI (*74, 75*) to classify ASD pathology [reviewed in (*21*)]. Our approach is the first to use a multiscale, ultra high-resolution dataset based on microscopy that can examine and bridge fine structural, molecular, histopathological, and functional pathway attributes to classify patterns of brain network/circuit organization in the neurotypical brain and in ASD. In future studies, we can use the ACC trained network to examine other brain regions that have or do not have known pathology in ASD. Accurate classification of images in a new region “B”, using the already trained ACC-DNN would suggest that the nature of neuropathology in the two regions (ACC and B) is likely consistent. On the other hand, incorrect classification of images in a new region “B”, using the already trained ACC-DNN, would suggest either that region “B” is not affected in ASD, or that the nature of neuropathology in the two regions (ACC and B) is different and region dependent. In the latter case, a new, optimized DNN, would need to be developed and trained with images from both ACC and any new region(s) to maximize performance and increase accuracy. This way, our proposed method can be generalized and applied to detect axonal and pathway feature differences across brain areas in the neurotypical brain and in psychiatric disorders with underlying pathology in brain network connectivity. This methodical approach holds great promise for the reliable implementation of DNN training and systematic testing to study many brain areas and networks, which would otherwise take prohibitively long to examine, and would provide key insights that will guide future studies and development of targeted diagnostics and interventions.

## Acknowledgments

This work was supported by a grant from the Simons Foundation (SFARI Grant 946867) to BZ and AY. Some tissues were obtained from Autism BrainNet, a resource of the Simons Foundation Autism Research Initiative (SFARI). Autism BrainNet also manages the Autism Tissue Program (ATP) collection, previously funded by Autism Speaks. We are grateful and indebted to the families who donated tissue for research purposes to Autism BrainNet and the ATP. We also gratefully acknowledge brain donors and their families, the Harvard Brain Tissue Resource Center, the Institute for Basic Research in Developmental Disabilities, the University of Maryland Brain and Tissue Bank, and the National Disease Research Interchange for providing post-mortem human brain tissue. We thank Tara McHugh, for technical assistance in cutting and staining the human specimens.

## Author contributions

Conceptualization: AY and BZ; Experimental procedures: BZ, XL, SL; Analysis: KD, KK, TL, AY, BZ; Funding acquisition: AY, BZ; Writing – original draft: KD, KK, AY, BZ; Writing – review & editing: AY, KD, KK, TL, XL, SL, BZ.

## Data availability

Data supporting the findings of this study are either included in the manuscript and supplementary materials or are available from the corresponding author upon request.

## Code availability

The code is deposited at https://github.com/kkuang0/ASD-ACC-Transfer-Net

