## Supplementary Figures for "Artificial intelligence networks combining histopathology and machine learning can extract axon pathology in autism spectrum disorder"

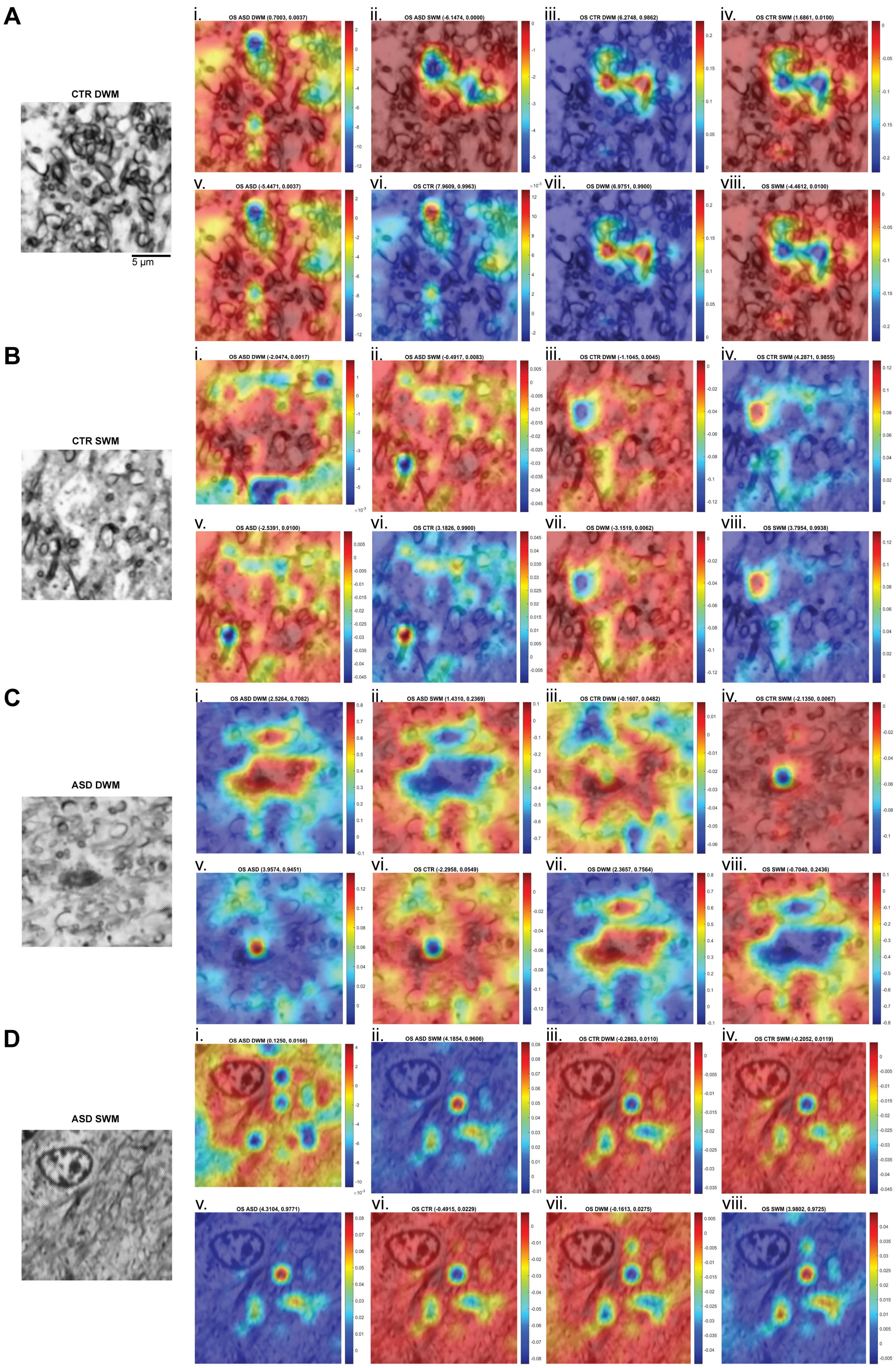
A
CTR DWM
5 μm
i. OS ASD DWM (0.7003, 0.0037)
ii. OS ASD SWM (-6.1474, 0.0000)
iii. OS CTR DWM (6.2748, 0.9862)
iv. OS CTR SWM (1.6861, 0.0100)
v. OS ASD (-5.4471, 0.0037)
vi. OS CTR (7.9609, 0.9963)
vii. OS DWM (6.9751, 0.9900)
viii. OS SWM (-4.4612, 0.0100)
B
CTR SWM
i. OS ASD DWM (-2.0474, 0.0017)
ii. OS ASD SWM (-0.4917, 0.0083)
iii. OS CTR DWM (-1.1045, 0.0045)
iv. OS CTR SWM (4.2871, 0.9855)
v. OS ASD (-2.5391, 0.0100)
vi. OS CTR (3.1826, 0.9900)
vii. OS DWM (-3.1519, 0.0062)
viii. OS SWM (3.7954, 0.9938)
C
ASD DWM
i. OS ASD DWM (2.5264, 0.7082)
ii. OS ASD SWM (1.4310, 0.2369)
iii. OS CTR DWM (-0.1607, 0.0482)
iv. OS CTR SWM (-2.1350, 0.0067)
v. OS ASD (3.9574, 0.9451)
vi. OS CTR (-2.2958, 0.0549)
vii. OS DWM (2.3657, 0.7564)
viii. OS SWM (-0.7040, 0.2436)
D
ASD SWM
i. OS ASD DWM (0.1250, 0.0166)
ii. OS ASD SWM (4.1854, 0.9606)
iii. OS CTR DWM (-0.2863, 0.0110)
iv. OS CTR SWM (-0.2052, 0.0119)
v. OS ASD (4.3104, 0.9771)
vi. OS CTR (-0.4915, 0.0229)
vii. OS DWM (-0.1613, 0.0275)
viii. OS SWM (3.9802, 0.9725)

**Supplementary Figure 1: Image sensitivity maps of correctly classified Tol-Blue microscopy images. A-D,** Representative examples of Tol-Blue images and overlaid heat maps from each of the 4 classes: CTR DWM (A), CTR SWM (B), ASD DWM (C), ASD SWM (D). For each class we used Occlusion Sensitivity to generate 8 heatmaps, representing hotspots and their relative weights (probability) for correct and incorrect class assignment: **i.** ASD DWM, **ii.** ASD SWM, **iii.** CTR DWM, **iv.** CTR SWM, **v.** ASD, **vi.** CTR, **vii.** DWM, **viii.** SWM. The title of each heatmap shows the sum of model neuron activity from the fully connected layer, as well as the SoftMax probability. Heatmaps of the correct corresponding class highlight areas of the image that contributed most towards the overall classification. Heatmaps of the other classes provide a comparative visualization of how different features were weighted. This aids in identifying specific patterns and features that distinguish each individual class, enhancing the interpretability and transparency of the network's performance.

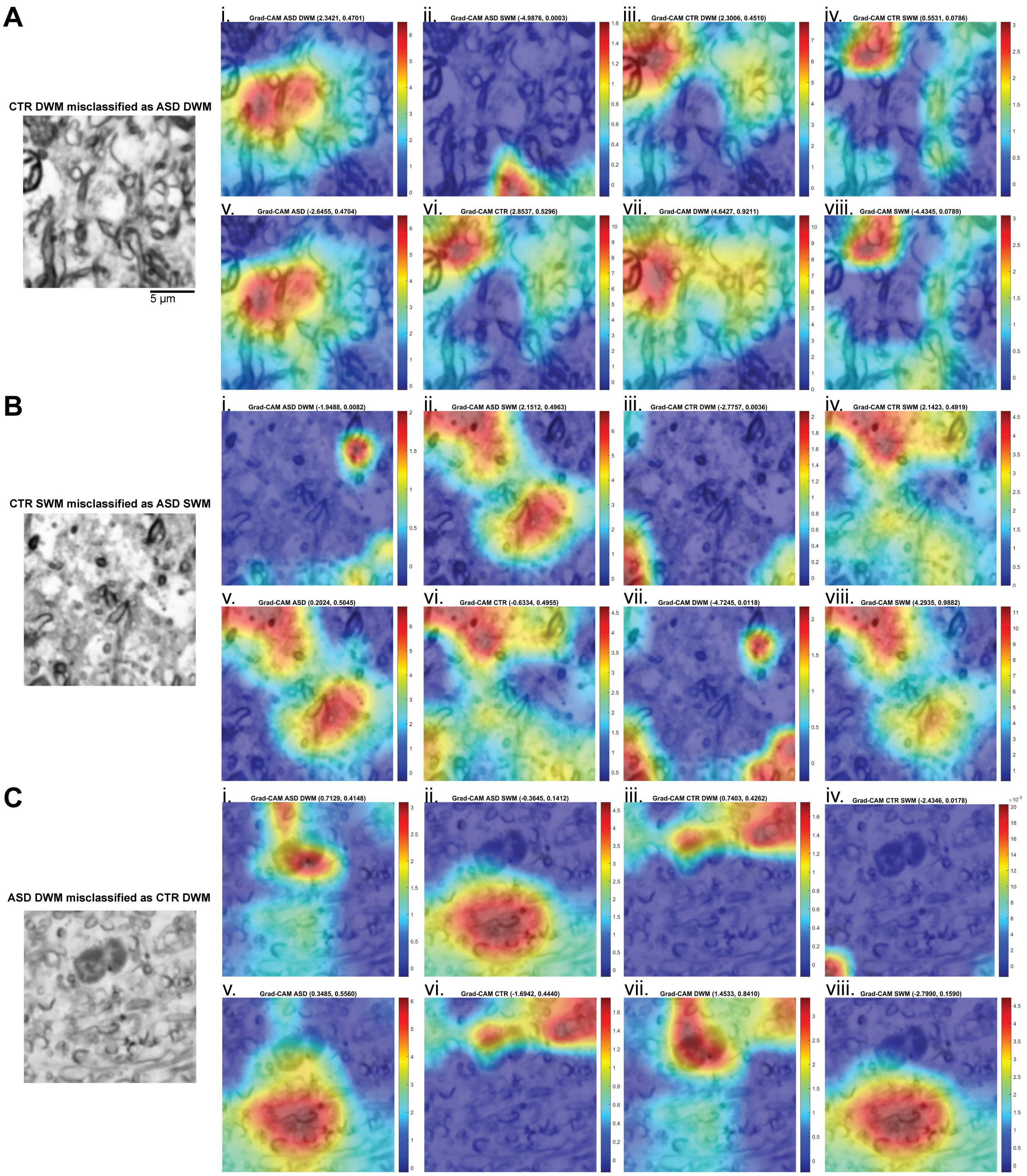


**Supplementary Figure 2: Image sensitivity maps of incorrectly classified Tol-Blue microscopy images. A-D,** Representative examples of Tol-Blue images and overlaid heat maps from each of the 4 classes: CTR DWM (A), CTR SWM (B), ASD DWM (C), ASD SWM (D). For each class we used Grad-CAM to generate 8 heatmaps, representing hotspots and their relative weights (probability) for correct and incorrect class assignment: **i.** ASD DWM, **ii.** ASD

SWM, **iii.** CTR DWM, **iv.** CTR SWM, **v.** ASD, **vi.** CTR, **vii.** DWM, **viii.** SWM. The title of each heatmap shows the sum of model neuron activity from the fully connected layer, as well as the SoftMax probability. Heatmaps of the predicted corresponding class highlight areas of the image that contributed most towards the overall classification. Heatmaps of the other classes provide a comparative visualization of how different features were weighted. This aids in identifying specific patterns and features that distinguish each individual class, enhancing the interpretability and transparency of the network's performance.
